# Three novel deep-branching orphan protist lineages from the marine rare biosphere

**DOI:** 10.64898/2026.11.16.766945

**Authors:** Nina Pohl, Miguel M. Sandin, Veera Tuovinen Nogerius, Fabien Burki

## Abstract

Metabarcoding has transformed our view of marine protist communities, yet many sequences in environmental surveys are not assigned confident taxonomic identity. These taxonomically ambiguous sequences may include lineages that are not within any known eukaryotic supergroup, which we refer to here as orphan lineages. These orphan lineages matter beyond taxonomy-based community analyses, because they can help us understand early eukaryotic evolution. Here we used phylogenetic inferences and sequence similarity networks (SSNs) of long-read metabarcoding data of the ribosomal operon (18S-28S rDNA) to identify three novel phylum-level orphan lineages. We designate these lineages Deep Orphan Lineages 1-3 (DEEPOLs) and show that they are exclusively marine and part of the rare biosphere, yet ecologically distinct: DEEPOL1 is a cosmopolitan generalist, DEEPOL2 a conditionally rare lineage capable of local polar blooms, and DEEPOL3 a non-polar lineage with a tendency toward the dark ocean. For DEEPOL2, lineage-specific digital PCR (dPCR) combined with catalyzed reporter deposition fluorescence in situ hybridization (CARD-FISH) provides the first in situ visualization of these cells, revealing small, flagellated, and morphologically inconspicuous protists. Together, our results show that substantial diversity remains hidden among rare lineages, and that long-read sequencing combined with targeted molecular and microscopy approaches offers a first step toward linking orphan sequences to organisms and their evolution and ecological roles.

## Introduction

The eukaryotic tree of life describes the evolutionary relationships among all eukaryotes, from the root of the tree to the largest eukaryotic clades known as supergroups. The tree has been profoundly reshaped in the last 20 years, importantly through the discovery of orphan lineages. These lineages are protist groups that do not belong to already established supergroups and thus represent some of the highest ranked eukaryotic assemblages. Protist orphan lineages occupy pivotal positions for reconstructing the broadscale evolutionary tree of eukaryotes [1]. They provide critical reference points for resolving deep relationships among eukaryotic supergroups and for inferring the evolutionary innovations that have shaped cellular organization, trophic strategies, and metabolic capacities [2–4]. Beyond their evolutionary relevance, protists, including orphan lineages, contribute substantially to the functional diversity of eukaryotes, most of which remains poorly characterized [5,6]. As integral components of microbial food webs, protists influence energy flow, nutrient recycling, and ecosystem stability [7].Yet these lineages have largely been discovered through low-throughput, laborious manual cell isolation and culturing, an approach restricted to organisms that can be found and grown. The vast body of cultivation-independent environmental sequence data, by contrast, has rarely yielded comparable discoveries.

Several notable protist lineages have been discovered or re-isolated in recent years. Some of these were orphans when they were first detected and have since been placed in the eukaryotic tree. These include *Meteora* and *Solarion*, which together with Hemimastigophora and Provora form the recently proposed supergroup Promethea (Disparia) [2,8], and Picozoa, which represent a major branch in Archaeplastida [9]. These findings highlight, if needed be, that the catalogue of major eukaryotic lineages known to science is not complete. Common features of these lineages are that they are heterotrophic, and often–but not always–rare and undetected in high-throughput environmental molecular surveys [10].

In recent years, however, high-throughput single-cell and shotgun metagenomic sequencing have shown that such lineages can be detected directly from environmental sequence data. Single-cell amplified genomes (SAGs) have revealed novel lineages with no clear affinities to existing supergroups [11], and have expanded the number of mitochondrial genomes from underexplored lineages such as Picozoa and Telonemia [12]. Organellar metagenomics has also uncovered novel eukaryotic diversity. The compact, gene-dense organization of organellar genomes resembles that of prokaryotic genomes, which makes them well suited to the genome-resolved metagenomic approaches widely applied to bacteria and archaea [13–17]. Examples include the newly described algal lineage Leptophytes, identified in a dataset of plastid metagenome-assembled genomes (MAGs) [18], and mitochondrial MAGs belonging to Cryptophyceae [19]. Even nuclear genomes, although harder to assemble than organellar ones, have been reconstructed from comprehensive metagenomic datasets such as Tara Oceans [20]. However, both single-cell genomics and metagenomics remain biased toward relatively abundant organisms, so rare diversity may escape detection by these approaches [20].

The method that typically covers most protist diversity in environmental surveys, including rare taxa, is metabarcoding. However, the relatively limited taxonomic resolution of standard short-read metabarcoding sequences, especially for groups with sparse or no reference sequences, translates into a substantial part of protist molecular diversity with undetermined origin. Even in the euphotic ocean, one of the best-sampled environments, approximately 20 % of protist sequences still cannot be taxonomically assigned. In less explored habitats, such as deep-sea sediments, this proportion reaches 60 % [21,22]. However, short-read metabarcoding has pitfalls that weigh most heavily in the rare biosphere. Common markers, such as the V4 and V9 regions of the 18S rRNA gene, offer limited phylogenetic resolution and rely on well-annotated reference sequences, which for much of the diversity do not exist [23,24], while PCR and sequencing errors, chimeras, and amplification biases can create or distort low-abundance sequences [25,26]. Orphan sequences should therefore be interpreted with care and supported by multiple lines of evidence, such as their recurrence across independent samples and studies, robust phylogenetic placement, and, ideally, visualization of the cells themselves. Longer molecular markers can strengthen several of these pitfalls at once. Sequencing large regions of the ribosomal operon (∼5 000 bp) from environmental DNA with long-read platforms such as Pacific Biosciences (PacBio) and Oxford Nanopore Technologies (ONT) have provided more phylogenetic information and potentially offers new opportunities to identify novel deep-diverging protist lineages [24,27–30].

Here, we used a long-read metabarcoding dataset spanning multiple environments [28] to detect previously unrecognized orphan lineages. Using a combined phylogenetic and SSN approach, we identify three robust clades composed exclusively of orphan sequences. We then integrate short-read metabarcoding data to characterize their biogeography worldwide and use lineage-specific dPCR and CARD-FISH combined with protein staining to visualize one lineage *in situ*, linking molecular novelty to cellular morphology.

## Material & Methods

### Identification of orphan clades from long-read metabarcoding data

The long-read metabarcoding dataset used for the identification of orphan lineages was available from [28]. To identify novel orphan lineages, we applied two complementary approaches to the long-read sequencing data. First, the published phylogenetic tree [28] was screened to identify monophyletic clades of orphan sequences, defined as sequences that were assigned to the domain Eukaryota and lacked affiliation with any described lineage at lower taxonomic ranks. Using a custom R script (https://github.com/burki-lab/deep-orphan-lineages) we retrieved the descendant tips of every internal node of the rooted tree and retained nodes in which all tips were orphan sequences and which contained at least two sequences. Where such clades were nested, only the largest one was kept. These clades, which contain no sequence from an established eukaryotic supergroup, were considered candidate orphan clades. Second, SSNs were constructed from the same sequences to independently assess sequence clustering. Pairwise similarities were calculated using all-against-all BLASTn, and networks were generated across a range of identity thresholds (≥80, ≥85, ≥87.5, ≥90, ≥92.5, ≥95, ≥97, and ≥99 %) with an additional minimum alignment coverage requirement of 40 % using python scripts (https://github.com/MiguelMSandin/random/tree/main/networks). Candidate lineages were identified using network analyses focusing on identifying maximal cliques (groups of sequences that are all connected to each other) composed exclusively of orphan sequences and assessing their similarity to annotated supergroups using shortest-path analyses. Candidate deep orphan lineages were retained only when supported by both approaches, such that the same set of sequences formed a monophyletic clade in the phylogeny and a corresponding maximal clique in the SSNs.

To verify that the newly identified DEEPOL lineages did not correspond to known lineages absent from our long-read metabarcoding dataset because they had not yet been described at the time of the study, DEEPOL sequences were queried against the NCBI nr/nt database using the web interface (last accessed October 2026) and locally against the Protist Ribosomal Reference database (PR2 v5.0.0) using BLASTn (v2.6.0+). All hits with ≥95 % sequence identity and an e-value of 0 were retained for phylogenetic validation, together with the 20 best-scoring hits of formally named species for each individual sequence within each DEEPOL lineage, to provide broader phylogenetic context. Duplicate hits shared between sequences were subsequently removed. DEEPOL sequences and the retained BLAST hits were placed in a phylogenetic tree covering the major eukaryotic supergroups, built from 18S rRNA gene sequences from the PR2 database. Sequences were aligned using MAFFT (v7.515) [31] with automatic parameter selection, followed by trimming with TrimAl (v1.4.rev15) [32] to remove positions containing more than 95 % gaps. Maximum-likelihood phylogenetic trees were inferred with IQ-TREE (v2.3.6) [33] using ModelFinder for model selection and 1000 ultrafast bootstrap replicates.

### Retrieval of DEEPOL-affiliated ASVs from short-read metabarcoding data

Long-read metabarcoding sequences carry higher phylogenetic signal for environmental diversity analysis than short-read barcodes, but their availability remains limited. To assess the diversity and distribution of the identified DEEPOL lineages, the DEEPOL 18S rRNA gene sequences were thus queried locally against the EukBank metabarcoding dataset, a curated collection of global V4 18S rRNA gene amplicon sequence variants [24], using BLASTn (v2.6.0+). Hits with ≥95 % identity and an e-value ≤1 × 10^-10^ were retained for phylogenetic validation. For each lineage, a tree was inferred as described above from all retained ASVs, the long-read DEEPOL sequences and selected Haptista sequences from PR2 as outgroup. ASVs forming a well-supported (bootstrap ≥95 %) monophyletic clade with the respective DEEPOL were retained for further analyses.

### Biogeographic analyses of DEEPOL lineages

The distribution of DEEPOL1-3 was assessed using metadata associated with the EukBank dataset [34]. For each lineage, amplicon sequence variant (ASV) read counts were divided by the total number of reads in the respective sample (sequencing depth) to calculate relative abundances, and ASVs detected in fewer than three samples were excluded. Community composition analyses were based on presence–absence matrices, which avoid the strong skew of metabarcoding read counts and reduce the influence of differences in sequencing depth, rRNA gene copy number and PCR amplification, and summarise lineage occurrence across habitats, ocean basins, and depth layers, with community dissimilarity calculated using Jaccard distance. Ocean basins were assigned by intersecting sample coordinates with IHO Sea Areas polygons (v3) [35]. Differences in lineage community composition were evaluated using permutational multivariate analysis of variance (PERMANOVA) with 999 permutations. Because PERMANOVA can be sensitive to differences in variability between groups, permutational analysis of multivariate dispersions (PERMDISP) was used to test whether significant results reflected differences in within-group dispersion (the spread of samples around their group centroid) rather than shifts in community composition. Multivariate analyses of community composition were applied separately to each DEEPOL lineage and included only marine samples in which the lineage was detected, with an assigned ocean basin, known sampling depth and a size fraction of pico- (0.2-3 µm), nano- (3-20 µm) or microplankton (20-200 µm). ASVs present in only one of these samples were removed. Samples were grouped by polar region (polar: Arctic Ocean or south of 60°S; non-polar: all other marine samples) and ocean layer (sunlit: sampled at ≤200 m; dark: >200 m). These filters did not apply to the coastal affinity analysis, which used a broader sample set (see below).

To test for coastal versus open-ocean preferences, the distance from each marine sample to the nearest coastline was calculated from its coordinates (Natural Earth; sf) [36], and samples within 200 km were classified as coastal. In contrast to the composition analyses above, this analysis included all marine samples with coordinates and sequencing depth, with missing or non-standard size fractions and depth layers coded as an additional category. Detection of each lineage was modelled with logistic regression, and relative abundance in positive samples (log10) with linear regression, using log10 distance to coast as predictor and polar region, ocean layer, size fraction and sequencing depth as covariates. ASV composition of coastal and open samples was compared with PERMANOVA and PERMDISP as above, and indicator ASVs were identified by indicator value analysis [37] with Benjamini-Hochberg correction.

All analyses were performed in R [38]. Core R packages included tidyverse (v. 2.0.0) [39] for data handling and manipulation, ggplot2 (v. 4.0.2) [40] for visualization, vegan (v. 2.7-3) [41] for distance-based community analyses, ordination, PERMANOVA, PERMDISP, distance-based redundancy analysis (dbRDA), and variation partitioning, ape (v. 5.8-1) [42] and ggtree (v. 3.16.3) [43] for phylogenetic handling and visualization, and UpSetR (v. 1.4.0) [44] for set-intersection analyses sf (v. 1.1.0) [36] and rnaturalearth (v. 1.2.0) [45] for distance to coast, broom (v. 1.0.12) [46] for model output, and indicspecies (v. 1.8.0) [37] for indicator analysis. Parts of the code for the analysis were debugged and restructured using AI assistance in the form of Claude Sonnet 5.5 (Anthropic). The code was reviewed, run and validated by the authors, who take full responsibility for the analyses and results.

### Design of DEEPOL2-specific 18S rRNA oligonucleotide

To increase the chances of successful detection, we focused subsequent probe design, dPCR, and CARD-FISH-based morphological screening on DEEPOL2, which reached the highest relative abundances in the EukBank dataset, whereas DEEPOL1 and DEEPOL3 were considerably less abundant.

A lineage-specific oligonucleotide targeting the 18S rRNA gene of DEEPOL2 was designed using the oligoN-design pipeline [47], with long-read DEEPOL2 metabarcoding sequences as targets and the PR2 database used for specificity assessment. Candidate oligonucleotides were evaluated based on sequence conservation, thermodynamic properties, predicted accessibility, and potential off-target matches, and the oligonucleotide achieving the highest combined sensitivity and specificity was selected. To further exclude unintended matches to non-target taxa not represented in PR2, additional BLASTn searches were performed against the NCBI nr/nt database.

The selected target site was used both as a lineage-specific CARD-FISH probe and as the forward primer in a dPCR assay. Reverse primers generated by the oligoN-design pipeline were screened for melting temperature, GC content, and predicted secondary structure. The reverse primer was selected to be located 68 bp downstream of the forward binding site, producing a 104 bp amplicon suitable for efficient amplification in dPCR. The dPCR primers deepol2_fwd (5′-TGAGAAAGGGGAATGGTT-3′) and deepol2_rev (5′-AAACGTTCTTGCCAAATG-3′) were synthesised as unmodified oligonucleotides by Sigma-Aldrich (Merck, Darmstadt, Germany). The CARD-FISH probe deepol2 (5′-AACCATTCCCCTTTCTCA-3′) was ordered from Biomers.net (Biomers, Ulm, Germany) with a 5′ horseradish peroxidase (HRP) modification.

### dPCR Screening of Samples for DEEPOL2

A total of 67 marine DNA extracts archived in our laboratory were screened for the presence of DEEPOL2. The samples originated from routine marine sampling conducted over several years at 12 locations (Table S11). Seawater was filtered onto track-etch polycarbonate membrane filters (25 mm diameter, 5 µm or 0.2 µm pore size; or 47 mm diameter, 1 µm pore size; Whatman, Cytiva, Little Chalfont, UK) using a peristaltic pump and Swinnex filter holders (Millipore, Merck KGaA, Darmstadt, Germany). Filters were flash-frozen in liquid nitrogen and stored at -80 °C until DNA extraction.

DNA was extracted using either the DNeasy PowerSoil Pro Kit or the DNeasy Plant Kit (Qiagen, Hilden, Germany), following the manufacturer’s instructions with minor modifications. For PowerSoil Pro extractions, samples underwent three freeze–thaw cycles (liquid nitrogen/65 °C), followed by bead beating (two 30 s cycles at 30 s^-1^ using a TissueLyser II and an additional 10 min vortex step). For Plant Kit extractions, filters were transferred to Precellys lysing tubes (Bertin Technologies, Montigny-le-Bretonneux, France) with AP1 buffer, subjected to three freeze–thaw cycles, bead beating for 2 min at 30 s⁻¹, and incubation with Proteinase K (45 µL, 55 °C, 1 h). DNA was eluted in 40 µL elution buffer and stored at −20 °C until analysis.

dPCR was performed on a QIAcuity system using the EG PCR Kit (Qiagen, Hilden, Germany). Reactions (40 µL) contained 0.5 µM of each primer (deepol2_fwd and deepol2_rev). Primer performance and assay sensitivity were verified using a synthetic gBlock containing the DEEPOL2 target region (10³ copies per reaction; Integrated DNA Technologies, Coralville, IA, USA; sequence in Table S12). Thermal cycling consisted of an initial denaturation at 95 °C for 2 min, followed by 40 cycles of 95 °C for 15 s, 54 °C for 30 s, and 72 °C for 30 s, with a final hold at 40 °C for 5 min. Fluorescence imaging was performed with an exposure time of 200 ms and a gain of 4. Thresholds were set manually for each sample based on fluorescence histograms, and samples were considered DEEPOL2-positive when a clearly separated population of positive partitions exceeded both the threshold and the signal observed in non-template controls.

### CARD-FISH optimisation and application for DEEPOL2

For *in situ* visualization of DEEPOL2 using CARD-FISH, samples from the Skagerrak region (North Sea) and the Bellingshausen Sea (Southern Ocean) were used (Table S13). Skagerrak samples were collected at multiple depths and time points as part of routine laboratory sampling. Seawater was fixed within one week of sampling with 3.7 % (v/v) formaldehyde for 1 h at room temperature, filtered onto 47 mm track-etch polycarbonate membrane filters (1 µm pore size; Whatman, Cytiva, Little Chalfont, UK), washed three times with 0.2 µm-filtered seawater, air-dried, and stored at -20 °C until further processing. The Southern Ocean sample was a small piece of an archived filter collected during the Tempano cruise in 2002 [48], provided by a collaborator (see Acknowledgements).

CARD-FISH was performed following [49] with minor modifications. In brief, filter sections were embedded in 0.1 % (w/v) low-melting agarose to minimise cell loss, incubated in 0.1 M HCl for 10 min to inactivate endogenous peroxidases, washed in PBS, rinsed in Milli-Q water, and air-dried. Hybridization was carried out overnight at 35 °C using 18 µL hybridization buffer and 2 µL of the HRP-labelled deepol2 probe (working solution 50 ng µL^-1^). Post-hybridization washing was followed by tyramide signal amplification using Alexa Fluor 488-labelled tyramide in the presence of H2O2 for 1 h at 46 °C in the dark. Filters were washed, rinsed, air-dried, and mounted using a DAPI-containing mounting medium. The mounting medium consisted of glycerol, Vectashield (Vector Laboratories, Inc., Newark, CA, USA), and PBS at a ratio of 10:2:1 (v/v/v) supplemented with DAPI at a final concentration of 5 µg/mL.

Hybridization stringency was optimised by testing a formamide gradient from 20 % to 60 % (v/v) in 10 % increments. For each formamide concentration, probe-hybridized filter pieces and corresponding no-probe controls were processed identically. Optimal hybridization stringency was defined as the lowest formamide concentration producing strong, specific DEEPOL2 signals with minimal background fluorescence and preserved cell morphology. A formamide concentration of 30 % was selected for all subsequent applications.

For whole-cell visualization, an NHS-ester protein stain conjugated to DyLight 560 (Dyomics, Jena, Germany) was applied prior to CARD-FISH. The NHS-ester was diluted 1:700 in 0.1 M sodium bicarbonate buffer, and filter pieces were incubated for 1.5 h at room temperature in the dark, followed by two PBS washes and air-drying before CARD-FISH processing. Each sample was processed with and without NHS-ester staining to assess potential staining artefacts. Where the available filter pieces were big enough, a no-probe control was included to evaluate background fluorescence and nonspecific tyramide deposition, and a universal eukaryotic probe (EUK1209R) as positive control.

All imaging was performed using an inverted Leica Stellaris 5 confocal laser scanning microscope equipped with an HC PL APO CS2 63×/1.40 oil immersion objective. Acquisition was done in sequential mode to avoid crosstalk between the channels. A 405 nm diode laser was used for DAPI (detection: 430-501 nm) and the tunable White Light Laser at 498 nm for Alexa Fluor 488 (503-582 nm), 578 nm for the NHS-ester DY560 (583-647 nm), and 649 nm for chlorophyll autofluorescence (654-783 nm). Z-stacks were acquired with a system-optimized step size of ∼0.30 µm, with total stack depth adjusted per sample to capture the signal from the whole cell. The Z-stacks are shown as maximum intensity projections (MIP), generated using Leica LAS X software. Brightness and contrast (LUT) were adjusted uniformly across compared images for display clarity.

## Results

### Identification and phylogenetic characterization of three novel orphan protist lineages

Phylogeny-guided taxonomic annotation of 16 821 long-read metabarcoding sequences were taken from [28]. Of these, 305 sequences were only annotated to the highest taxonomic rank in the analysis, i.e. domain Eukaryota, thus putatively representing orphan sequences. Here, we define an orphan lineage using two independent approaches: as a distinct clade branching outside all major eukaryotic lineages in phylogenetic trees, and simultaneously as a maximal clique (groups of sequences that are all connected to each other) in sequence similarity networks (SSNs).

Three lineages met these two criteria. We refer to them as Deep Orphan Lineages 1-3 (DEEPOL1-3), and together they comprise 14 of the 305 orphan sequences (Fig. 1). Because several deep-branching groups, including Provora, *Meteora* and *Solarion*, have been described since the long-read dataset was annotated [2,8,50], we next tested whether the DEEPOLs correspond to any of them. BLAST searches against the NCBI nucleotide database showed that all hits with ≥95 % identity were unnamed environmental sequences. No hits were assigned to a formally described species, and when added to our dataset, all formed well-supported groups with the respective DEEPOL, so we treated them as members of that lineage (Fig. S1). In contrast, similarity to named lineages was low (≤90 %), and phylogenetic analysis that included the recently described groups confirmed that the DEEPOLs represent novel uncharacterized diversity (Fig. S1). The remaining 291 orphan sequences were either placed within established eukaryotic supergroups or not recovered congruently by both approaches (Table S1).

**Figure 1.**
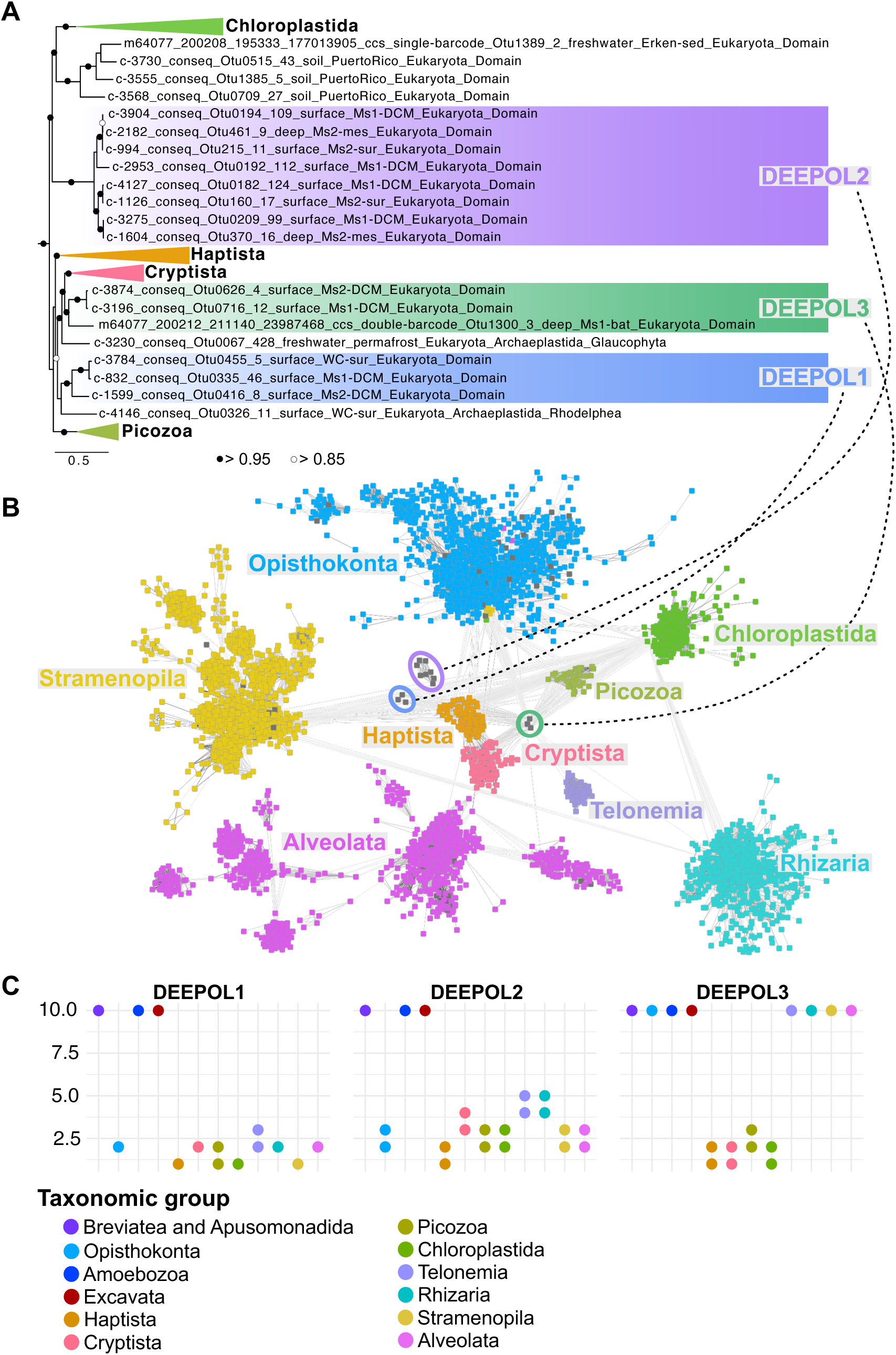
Identification of three deep-diverging protist lineages (DEEPOL1-3) using ribosomal phylogeny and sequence similarity networks. (**A**) Phylogenetic placement of DEEPOL1-3 inferred from long-read ribosomal operon sequences (modified from [28]). The three DEEPOL lineages form well-supported clades (>0.95 transfer bootstrap expectation (TBE)) distinct from established eukaryotic groups. (**B**) Sequence similarity network (SSN) constructed from the same long-read sequences, highlighting maximal cliques corresponding to the DEEPOL lineages. (**C**) Shortest-path analyses illustrating the graph-based distance between each DEEPOL clique and other annotated cliques in the SSN, shown from left to right for DEEPOL1, DEEPOL2 and DEEPOL3. Together, the congruence between phylogenetic inference and SSN structure supports the interpretation of DEEPOL1-3 as coherent deep-diverging evolutionary units.

In the phylogeny, the closest relatives of the DEEPOLs were plastid-bearing eukaryotic lineages or their close non-photosynthetic relatives, including Chloroplastida, Cryptista, Haptista, Stramenopila, Rhodelphis and Picozoa. In the SSN, the DEEPOLs were most closely connected to sequences from similar groups, although the specific groups differed slightly between the two approaches. For DEEPOL1, an alternative placement in the supergroup Promethea is also possible in the phylogeny (Fig. S1).

### Global distribution, habitat association, and size-fraction occurrence of DEEPOL lineages

To assess the global distribution, habitat association and size-fraction occurrence of the DEEPOLs, we retrieved corresponding ASVs from the short-read 18S rDNA V4 EukBank dataset and its metadata [34]. ASVs were retained only if they formed a well-supported monophyletic clade with the respective long-read DEEPOL sequences (Fig. S2-4). In total, 29 ASVs were assigned to DEEPOL1, 45 to DEEPOL2 and 12 to DEEPOL3. The mean sequence identities to the long-read DEEPOL sequences were 97.7 %, 98.3 % and 96.3 %, respectively, forming the basis of our biogeographic analyses (Table S3). Because the short V4 fragment carries little phylogenetic signal for deep eukaryotic relationships, some ASVs may be misassigned. However, given the well-supported phylogenetic placement and the consistency of distribution patterns (see below), we treat all retained ASVs as DEEPOL representatives.

All three DEEPOL lineages exhibit broadly cosmopolitan distributions but occur at low relative abundances. DEEPOL1 was consistently rare (median 0.009 %, SD = 0.03 %) and occurred across most ocean basins (Fig. 2A). DEEPOL2 showed a more heterogeneous and patchy distribution and was the most abundant lineage overall (median 0.03 %, SD = 0.3 %), with pronounced local maxima where relative abundance reached up to 5.3 % and exceeded 1 % in 58 samples, predominantly in the Southern Ocean (Fig. 2B). DEEPOL3 was the rarest and most sparsely distributed lineage (median 0.005 %, SD = 0.06 %). It was largely absent from polar regions and showed predominantly sporadic occurrence, but one sample from the South Pacific exhibited a high relative abundance of 2.03 % (Fig. 2C). Detection of the DEEPOLs was positively associated with sequencing depth, as expected for rare taxa, but absences were not restricted to shallowly sequenced samples: each lineage was also undetected in many deeply sequenced samples (Fig. S5). The lowest sequencing depths at which the lineages were detected were 769, 3 911 and 13 402 reads for DEEPOL1, DEEPOL2 and DEEPOL3, respectively.

**Figure 2.**
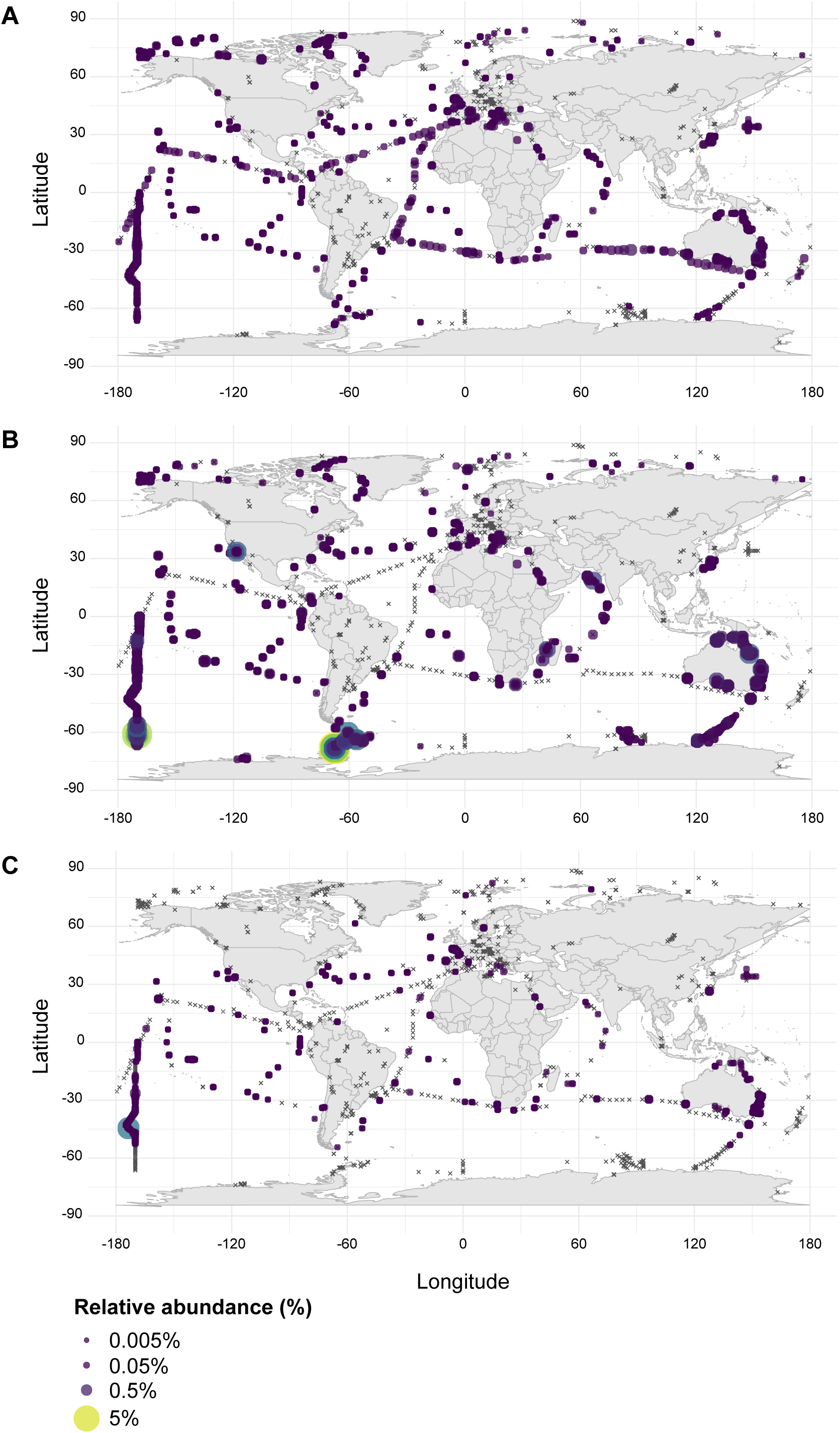
Global distribution and relative abundance of the DEEPOLs as found in the samples in the EukBank short-read 18S rRNA metabarcoding dataset. Maps are shown for DEEPOL1 (**A**), DEEPOL2 (**B**), and DEEPOL3 (**C**) separately. Dots indicate presence of the respective clade while crosses mark their absence. The size of the dots correspond to the average relative abundance of the DEEPOLs in the respective sample.

To test whether the DEEPOLs demonstrated a preference for coastal or open-ocean waters, we related detection and relative abundance to the distance to the nearest coast (Fig. 3). DEEPOL1 and DEEPOL2 were more likely to be detected farther from the coast (odds ratios per 10-fold increase in distance of 1.39 [95 % CI, 1.30-1.48], and 1.42 [95 % CI, 1.32-1.52], respectively; Table S4), whereas detection of DEEPOL3 did not vary with distance (1.05 [95 % CI, 0.97-1.14]). Among samples in which DEEPOLs were detected, relative abundance was unrelated to distance for DEEPOL1, about 11 % higher with every 10-fold increase in distance for DEEPOL2, and about 14 % lower for DEEPOL3 (Table S5). The effect of distance depended on ocean layer for DEEPOL1 and DEEPOL2 (interaction p < 10^-6^) and, for DEEPOL2, on polar region (p < 10^-8^; Table S7). In sunlit waters both lineages were detected more often offshore (odds ratios 1.39 [95 % CI 1.29-1.51], and 1.70 [95 % CI 1.54-1.88]), whereas in the dark ocean detection did not differ with distance, and abundance was higher closer to the coast (Table S6). Within polar waters, DEEPOL2 showed the opposite pattern, i.e. it was detected more often near the coast (odds ratio 0.63 [95 % CI 0.52-0.77]) than in non-polar waters (odds ratio 1.47 [95 % CI 1.35-1.60]). ASV composition differed between coastal and open-ocean samples in all three lineages (PERMANOVA, p ≤ 0.001), but explained little variation (R² = 0.3 to 2.7 %; Table S8). Indicator analysis identified mostly open-ocean ASVs: 10 of 29 in DEEPOL1, 6 of 45 in DEEPOL2 and 4 of 12 in DEEPOL3, against a single coastal indicator in DEEPOL1 and DEEPOL2 and none in DEEPOL3. Open-ocean indicators were largely consistent across ocean layers and polar niches, whereas the coastal indicators were not, and the DEEPOL1 coastal indicator ASV was significant in polar but not in non-polar waters (Table S9).

**Figure 3.**
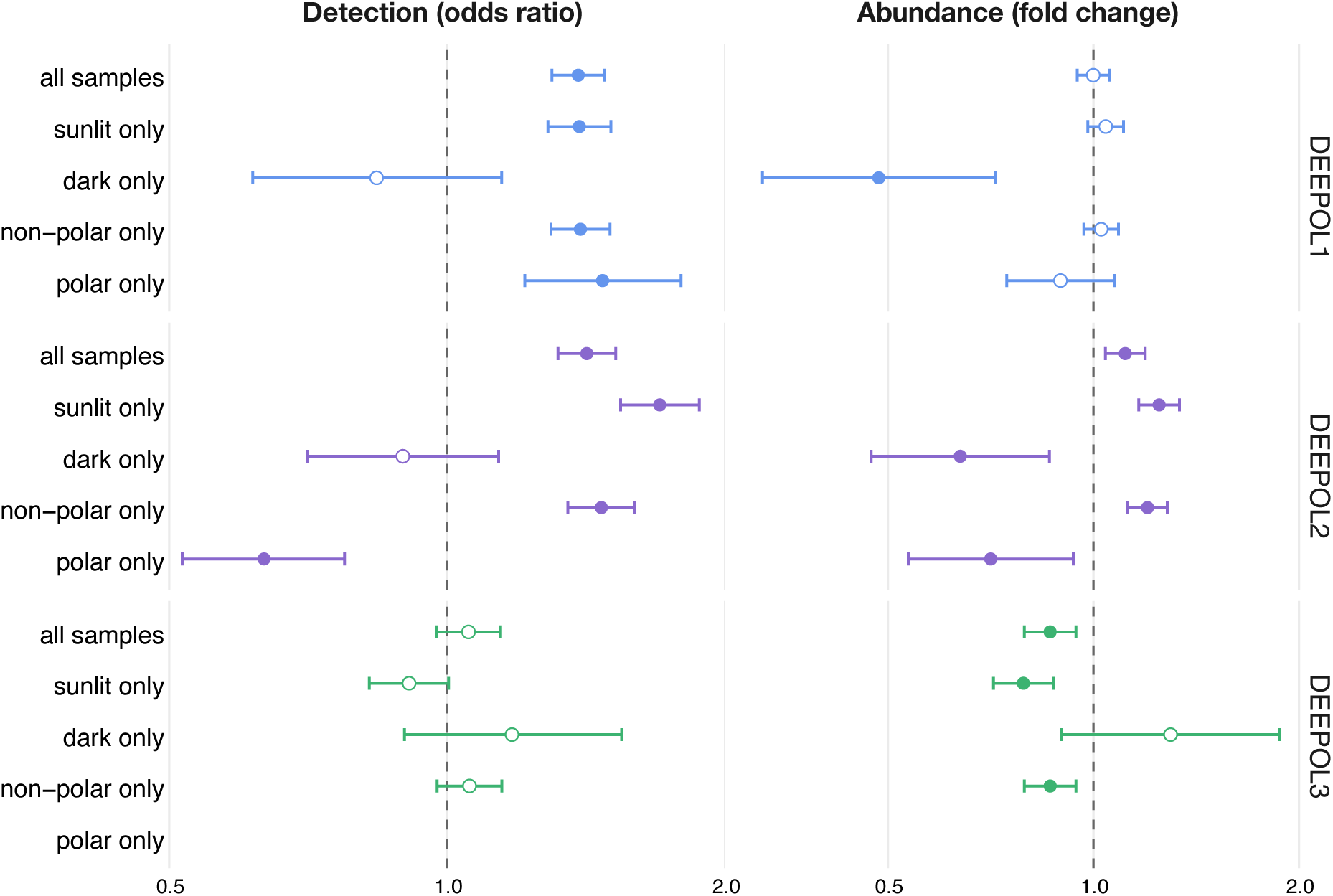
Association of DEEPOL detection and relative abundance with distance to the coast. Effect of a 10-fold increase in distance to the nearest coastline on the probability of detecting each lineage (left, odds ratio from logistic regression on all marine samples) and on its relative abundance in DEEPOL positive samples (right, fold change from linear regression of log10 relative abundance), shown for DEEPOL1, DEEPOL2 and DEEPOL3. Values below 1 indicate higher detection or abundance closer to the coast, and values above 1 indicate higher detection or abundance farther offshore. Points show estimates with 95% confidence intervals for all samples and within strata (sunlit, dark, non-polar and polar waters). Filled points indicate p < 0.05 and open points p ≥ 0.05 (unadjusted p-values).

Across the EukBank dataset, all DEEPOL lineages were strongly associated with marine environments, with only isolated detections in freshwater or soil samples (Fig. S6). DEEPOL1 was detected in 3 062 marine samples (31.3 % of marine samples), with a single terrestrial detection and no freshwater occurrences (Fig. S6A). DEEPOL2 exhibited the broadest habitat range but remained almost exclusively marine-associated, occurring in 3 554 marine samples (36.3 %), with rare detections in terrestrial (two samples) and freshwater systems (three samples; Fig. S6B). DEEPOL3 showed the most restricted habitat distribution, occurring exclusively in marine samples (1 359 samples; 13.9 %) and being entirely absent from terrestrial and freshwater datasets (Fig. S6C).

Most EukBank samples were generated using non-standard size fractionation or bulk filtration schemes, with only a subset employing strict pico- (0.2-2 µm), nano-(2-20 µm), or microplankton (20-200 µm) size fractionation. All three DEEPOL lineages were detected across multiple filtration strategies, but predominantly occurred in bulk and small size-fraction samples (Fig. S7A-C). To enable robust comparisons across standardized size classes, subsequent analyses were restricted to marine samples with well-annotated pico-, nano-, or microplankton fractions and complete depth and basin metadata. This filtering resulted in 401 samples and 28 ASVs for DEEPOL1, 301 samples and 36 ASVs for DEEPOL2, and 141 samples and 11 ASVs for DEEPOL3. DEEPOL 1-3 ASVs were primarily shared between pico-and nano-fractions, with minimal representation in the larger microplankton fraction (Fig. S7D-F). DEEPOL2 showed the greatest size-fraction overlap, including ASVs detected across all three fractions, whereas DEEPOL1 and DEEPOL3 exhibited more restricted distributions and lacked ASVs occurring exclusively in the microplankton fraction. Together, these patterns indicate that DEEPOL lineages are predominantly associated with small planktonic size classes, with substantial ASV-level overlap between pico-and nano-plankton.

### Environmental drivers of DEEPOL distribution

To assess large-scale structuring of DEEPOL occurrence across marine environments, multivariate analyses of presence–absence data based on Jaccard dissimilarities were performed. Principal coordinates analysis (PCoA) revealed lineage-specific structuring patterns: DEEPOL1 and DEEPOL2 were primarily organized along latitudinal gradients separating polar and non-polar samples, whereas DEEPOL3 was structured predominantly by water-column layer, with clear separation between sunlit and dark ocean samples (Fig. S8A-C). These patterns were supported by PERMANOVA, which identified polar versus non-polar classification as the dominant factor for DEEPOL1 (R² = 0.1433, p = 0.001) and DEEPOL2 (R² = 0.1501, p = 0.001), with depth contributing a smaller but significant fraction in both cases. In contrast, DEEPOL3 was primarily structured by depth layer (R² = 0.2056, p = 0.001), while latitudinal differentiation explained only a minor fraction of variance (R² = 0.0171, p = 0.001). PERMDISP analyses showed that dispersion contributed to some of the patterns specifically in DEEPOL1 and 2 (Fig. S8D-F, Table S4).

To identify the environmental variables underlying these lineage-specific patterns, dbRDA was applied using temperature, salinity, and depth as explanatory variables. For DEEPOL1, all three variables contributed significantly, with depth showing the strongest association, followed by temperature and salinity (Fig. 4A). DEEPOL2 showed a stronger environmental signal overall, with temperature emerging as the dominant driver, followed by depth and salinity (Fig. 4B). In contrast, DEEPOL3 remained primarily structured by depth, which explained the largest fraction of variation, exceeding the contributions of temperature and salinity (Fig. 4C). Variation partitioning across all three lineages indicated that environmental variables consistently explained a substantially larger proportion of variation than geographic factors, with relatively limited shared effects, emphasizing the dominant role of environmental gradients in structuring DEEPOL distributions (Table S10).

**Figure 4.**
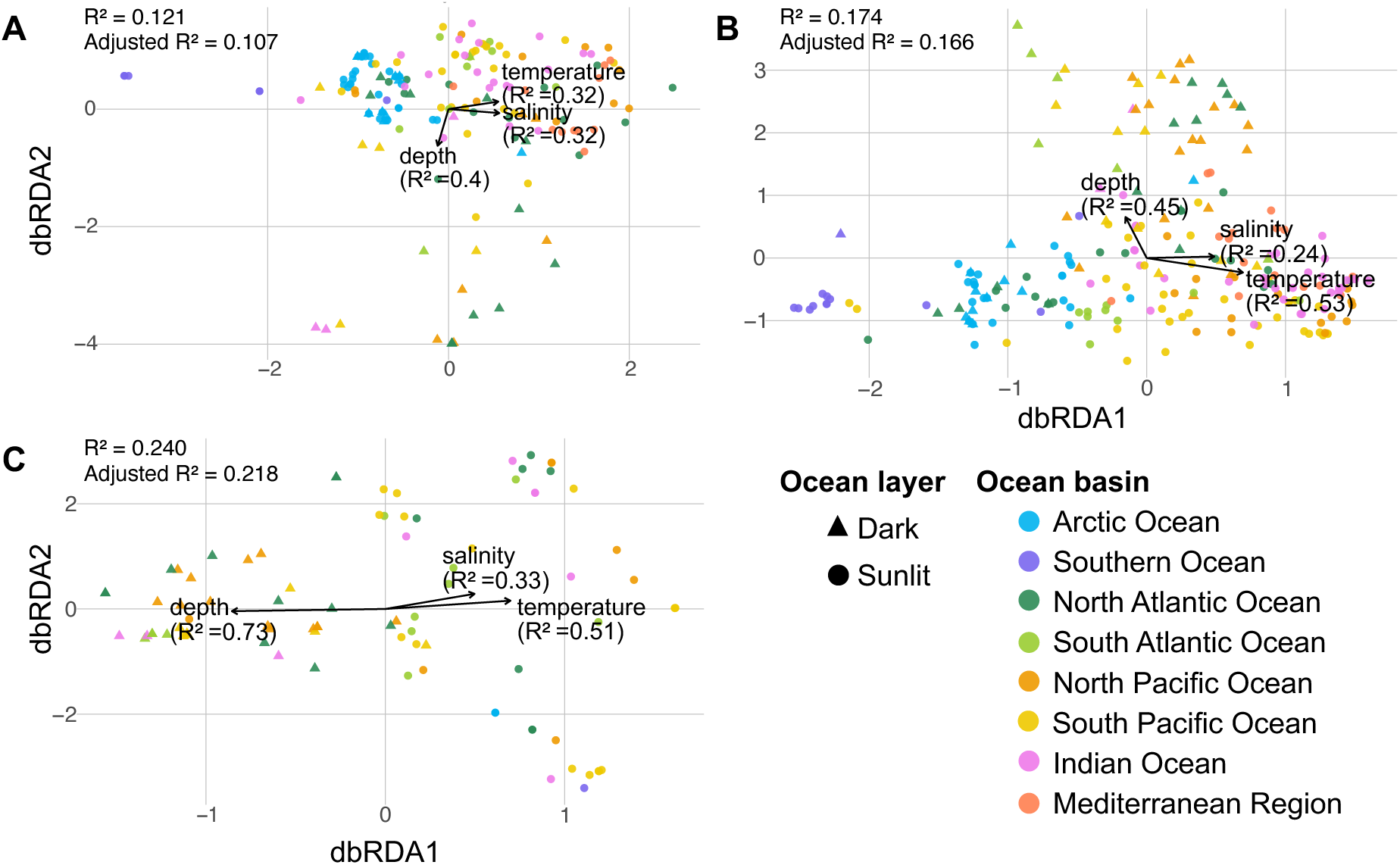
Distance-based redundancy analysis (dbRDA) of ASV presence/absence (Jaccard distance), illustrating the influence of significant environmental variables (envfit, p < 0.05) on lineage-specific community composition for DEEPOL1 (**A**), DEEPOL2 (**B**) and DEEPOL3 (**C**). The models were conditioned on latitude and longitude, so they show the effect of the environment independent of geographic position. Arrows indicate the fitted environmental variables, with the R² of each variable’s fit to the ordination in brackets. The R² and adjusted R² in each panel refer to the full model (temperature, salinity and depth, after conditioning on latitude and longitude). Samples from the sunlit ocean layers (0 to 200 m) are shown as circles, and samples from the dark ocean (>200 m) as triangles. Colours indicate the ocean basin each sample was taken from.

### Phylogenetic context of biogeography and ecological specialization in DEEPOL lineages

To assess whether large-scale biogeographic patterns and ecological specialization correspond to phylogenetic structure within DEEPOL lineages, ASV-level habitat associations were mapped onto lineage-specific phylogenetic trees. ASVs were classified as specialists when detected in ≥70 % of samples from a given oceanic layer (sunlit vs. dark) or latitudinal category (polar vs. non-polar), and all remaining ASVs considered generalists. ASV occupancy was quantified as the percentage of DEEPOL-positive marine samples in which each ASV was detected. Indicator ASVs of coastal (≤200 km from the coast) and open-ocean waters, identified across all samples (see above), were additionally displayed.

Across all lineages, ecological specialization differed between latitudinal and vertical habitat axes (Fig. 5A-C). DEEPOL1 and DEEPOL2 were dominated by non-polar specialists and by ASVs associated with sunlit-ocean samples, whereas DEEPOL3 consisted exclusively of non-polar specialists and showed a more balanced representation of sunlit-and dark-ocean specialists together with a small number of generalists. Phylogenetic resolution differed among lineages. In DEEPOL1, two subclades were consistently recovered with high support, one of which comprised predominantly sunlit-ocean, non-polar ASVs with high occupancy, while two polar-specialist ASVs were distributed across the larger clade, consistent with multiple independent transitions to polar environments (Fig. 5A). DEEPOL2 showed uneven phylogenetic resolution, with three of eight putative subclades well supported (bootstrap >95), whereas the deeper nodes remained unresolved (Fig. 5B; Fig. S3). All polar-associated ASVs were located in the largest subclade, which, however, was poorly supported (Fig. 5B). ASVs with high occupancy were distributed across multiple subclades and were all associated with sunlit-ocean conditions. In DEEPOL3, specialization to sunlit environments was the minority suggesting an overall preference for the dark ocean for this lineage. With one exception, all sunlit specialized ASVs were concentrated in one clade while the other two smaller clades were dominated by dark ocean specialists. The phylogeny of DEEPOL3 was further strongly structured by occupancy, with the most prevalent ASVs clustering within a single subclade, whereas remaining subclades were composed primarily of ASVs with lower detection frequencies (Fig. 5C). The two ASVs with highest occupancy were each specialized to the sunlit and the dark ocean, respectively, indicating that the lineage can be prevalent in either environment. Coastal and open-ocean indicator ASVs were few and scattered across the trees, without a clear phylogenetic pattern (Fig. 5A-C).

**Figure 5.**
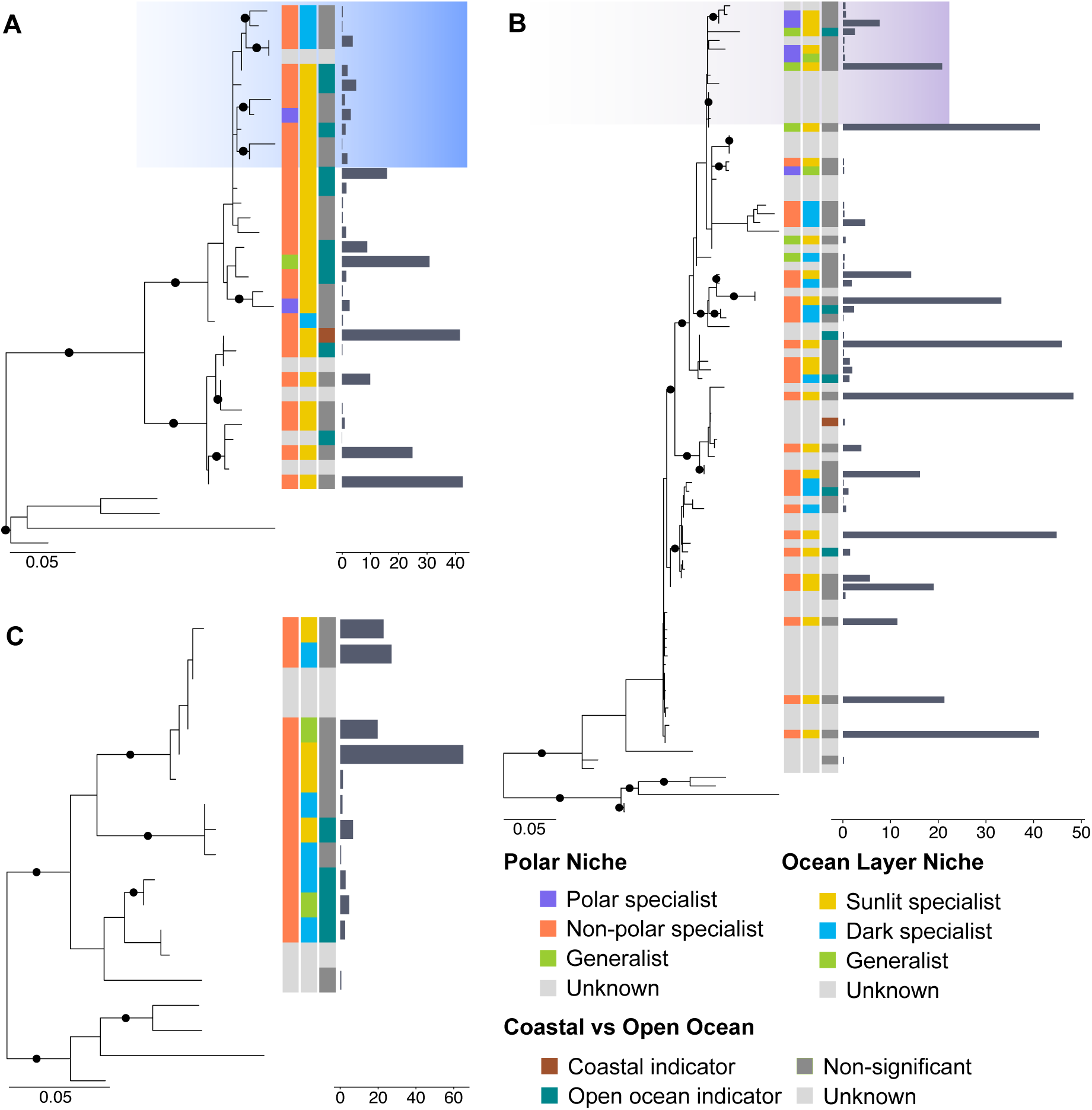
Lineage-specific 18S rRNA gene phylogenies of EukBank ASVs corresponding to DEEPOL1 (**A**), DEEPOL2 (**B**) and DEEPOL3 (**C**). For each ASV, the panels show (from left to right) its specialization for polar or non-polar waters, its specialization for the sunlit (0 to 200 m) or dark (>200 m) ocean layer, whether it is an indicator ASV of coastal waters (≤200 km from the coast) or of the open ocean (>200 km), and its occupancy, i.e. the percentage of DEEPOL-positive marine samples in which the ASV was detected. An ASV was classified as a specialist when at least 70% of its samples came from one category, and as a generalist otherwise. Coastal and open-ocean indicators were identified with indicator value analysis (IndVal.g, Benjamini-Hochberg adjusted p < 0.05); ASVs without a significant association are shown as not significant. Black circles indicate ultrafast bootstrap support ≥ 95%.

### Detection and *in situ* visualization of DEEPOL2 using dPCR and CARD-FISH

Metabarcoding has transformed our ability to assess the diversity of environmental microbial communities, yet the vast majority of cells behind the sequence tags have remained unseen. To link metabarcodes to cell morphology, we focused on DEEPOL2, which exceeded 1 % relative abundance at several locations (Fig. 2B) and was therefore the most promising candidate for visualization. To identify suitable samples, we screened 67 marine samples from four seas with a newly developed lineage-specific dPCR assay (Table S5). In line with the metabarcoding data, DEEPOL2 was detected in all regions but generally at low concentrations (mean = 15 copies L^-1^), however reaching a maximum of 258 copies L^-1^ (Fig. S9). We selected the three Skagerrak (North Sea) time-series samples with the highest DEEPOL2 concentrations for CARD-FISH, together with one Southern Ocean sample from the Bellingshausen Sea, where biogeographic analyses had predicted relatively elevated abundance (Fig. 2B). To characterize DEEPOL2 cell morphology in more detail, we combined specific probe fluorescence with DAPI and NHS-ester protein staining.

The lineage-specific probe deepol2 was predicted *in silico* to be highly specific to DEEPOL 2 as a whole. It matched all 18 DEEPOL2 sequences used for its design and had no perfect-match hits outside the lineage. Only two non-target organisms matched, *Trypanosoma* sp. and a species of Dinophyceae, with one and two mismatches, respectively. CARD-FISH successfully visualized DEEPOL2 cells in samples from both Skagerrak and the Bellingshausen Sea (Fig. 6; Fig. S10-12). Across all samples, the probe consistently hybridized to cells with similar characteristic morphology, further supporting the specificity of the hybridization. Negative controls, performed without the deepol2 probe, showed no fluorescence signal with a defined, cell-like morphology, confirming that the fluorescence observed in probe-treated samples reflects specific hybridization rather than autofluorescence or non-specific background staining (Fig. S13B, S14B, S15B).

**Figure 6.**
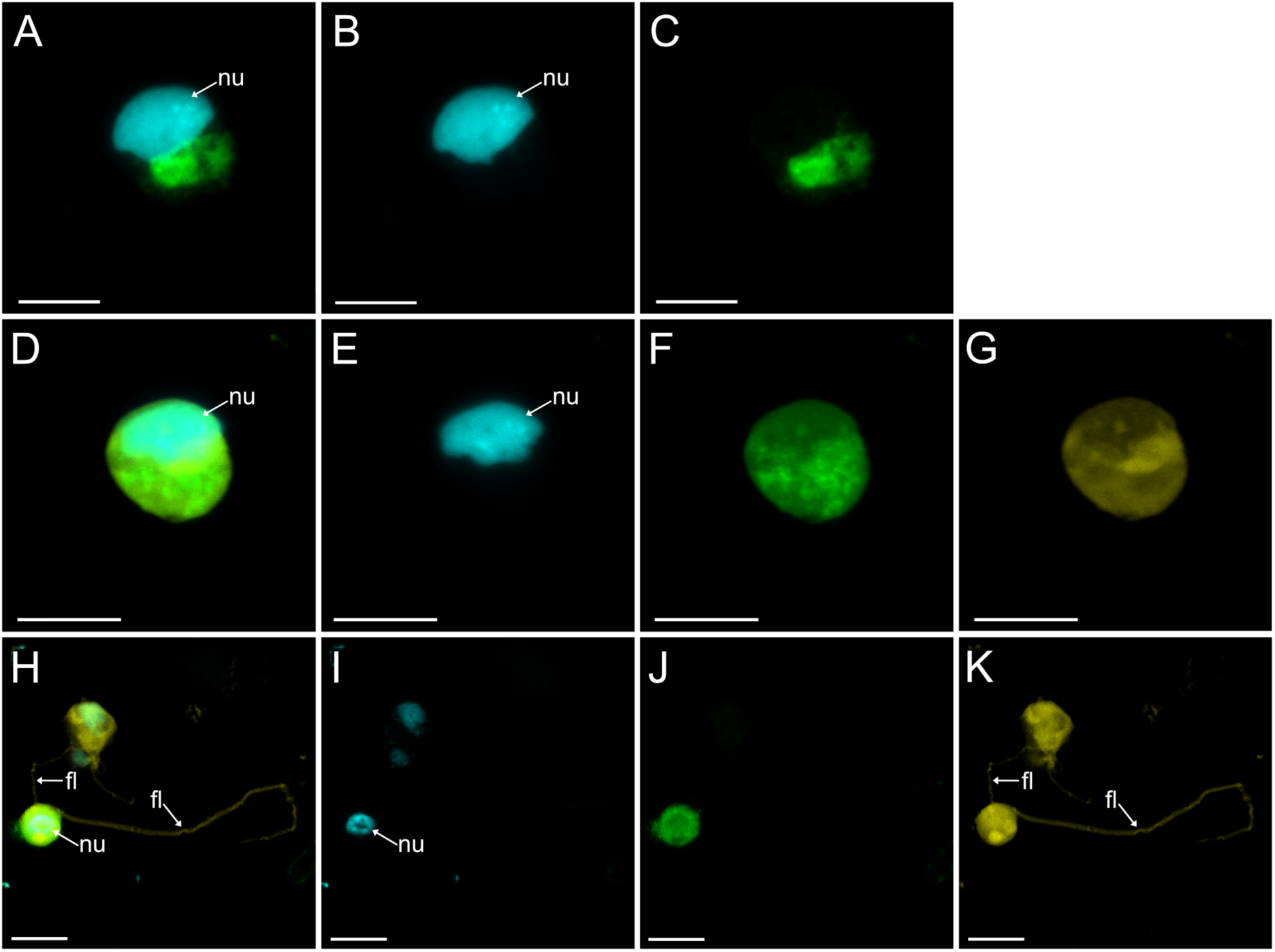
*In situ* visualization of DEEPOL2 cells using lineage-specific CARD-FISH. CARD-FISH was combined with NHS-ester protein staining to visualise DEEPOL2 cells in environmental samples. Panels show the overlay of all fluorescence channels (**A, D, H**), DAPI-stained nucleic acidsF, nu=nuclei (**B, E, I**), DEEPOL2-specific CARD-FISH probe signal detected with Alexa Fluor 488 (**C, F, J**), and NHS-ester protein staining using DY-560, fl=flagella (**G, K**). Panels **A-C** and **D-G** show DEEPOL2 cells from the North Sea (Skagerrak region), whereas panels **H-K** show a DEEPOL2 cell from the Southern Ocean (Bellingshausen Sea). Images shown represent maximum-intensity projections of Z-stacks acquired for each cell. Scale bars correspond to 5 µm.

The CARD-FISH signal was consistently bright relative to background fluorescence and spatially confined, often localized beneath or adjacent to the nucleus in cells without NHS-ester staining (Fig. 6A-C, Fig. S10, S11). The NHS-ester staining provided a clear representation of the outer cellular shape and flagella, but at the expense of a more diffuse and weaker probe signal than in cells without NHS-ester, possibly due to weak binding of the Alexa 488 to the NHS-ester (Fig. 6D-K; Fig. S10, S12). Across all samples, DEEPOL2 cells were spherical, with an average size of 5.1 × 4.5 µm (Table S14). Cells from the Skagerrak were slightly larger (mean 6.5 × 5.9 µm, length × width, n = 5) than those from the Bellingshausen Sea (mean 4.5 × 3.8 µm, n = 10; Table S14). NHS-ester-stained cells were smaller (mean 3.5 × 3.7 µm, n = 7) than cells without NHS-ester staining (mean 6.5 × 5.2 µm, n = 8), which may reflect shrinkage during the staining procedure, as incubation in sodium bicarbonate can affect cell volume. Prominent nuclei were observed in all cells (Fig. 6; Fig. S10-12). Flagella were consistently observed in Bellingshausen Sea DEEPOL2 cells, with two anterior flagella of unequal length, whereas no flagella were visible in the limited number of NHS-stained cells imaged from the Skagerrak samples (Fig. 6D-K; Fig. S10, S12). It remains unclear whether this reflects true morphological variation or methodological effects, such as breaking of flagella during the process of filtration.

## Discussion

In this study, we identified three previously unrecognized deep-branching protist lineages (DEEPOL1-3) from long-read metabarcoding sequences and provided biogeographic and morphological insights. Our results show that long-read metabarcoding provides sufficient phylogenetic signal to identify at least some deep-branching protist lineages. However, ribosomal genes-based phylogenies remain sensitive to challenges associated with deep evolutionary timescales, including signal erosion and long-branch artefacts [51–53]. Here, we used SSNs as a complementary, alignment-independent method for identifying these new orphan lineages (Fig. 1) [54,55]. The strong congruence between the deep-branching lineages recovered in both phylogenetic analyses and SSNs provides more robust evidence that the DEEPOLs are real lineages rather than artefacts of poorly resolved phylogenetic inference. The recovery of these DEEPOL lineages from a limited number of samples suggests that additional currently undetected deep-diverging eukaryotic diversity remains to be discovered, including many that are hidden in the high proportion of taxonomically unassigned sequences in short-read metabarcoding surveys [21,56,57]. Deep-branching lineages such as the DEEPOLs may therefore represent a persistent, but still sparsely sampled and almost totally uncharacterised, component of microbial eukaryotic diversity.

An important objective of this study was not only to identify orphan lineages but also to explore their biogeography and morphology in the absence of cultures or genomic data. Combining environmental metadata from large-scale short-read metabarcoding surveys with targeted *in situ* observations of DEEPOL2 places the lineages in a first ecological context. Across datasets, the DEEPOL lineages appear to be almost exclusively marine and consistently rare in relative abundance, with an average below 0.1 %, and occur across all ocean basins and a broad latitudinal range (Fig. 2). Such patterns are consistent with those typically observed for members of the rare microbial biosphere, a component of microbial communities increasingly recognized for its potential ecological significance despite low average abundance [10,58]. Rare lineages are thought to persist at low population sizes over long periods and may contribute to ecosystem functioning through condition-dependent activity or specialized ecological roles [59,60]. One example is *Symbiodinium*, the photosynthetic endosymbiont of reef-building corals: in the open ocean it accounts for only about 0.1 % of eukaryotic reads, yet it occurs at most stations, up to 3 000 km from reefs, and is transcriptionally active [61].

Low abundance alone does not explain why the DEEPOLs escaped detection for so long. Another important reason is that their inconspicuous morphology offers no clue to their phylogenetic position. For DEEPOL2 at least we can confirm this based on our CARD-FISH observations: the cells are small (2.0-7.7 × 2.5-6.9 µm, length × width) flagellates that resemble similar cells in different supergroups, for example *Nibbleromonas* that belongs to Provora (Fig. 6) [50], or *Palpitomonas* [62]. Such cells with superficially unremarkable morphology often do not attract attention in a light-microscopic survey, as their morphology offers no indication of their deep evolutionary divergence [63]. The lack of distinctive morphological feature has similarly delayed the recognition of other major environmental groups, including the MAST lineages, which remain known almost exclusively from sequence data despite their ecological prominence [64–66]. The morphological resemblance between DEEPOL2 and cells belonging to other supergroups indicates that the extent of undiscovered diversity is hard to assess. It is thus possible that a large overlooked uncharacterized diversity resides within the rare protist biosphere, owing to both low abundance and a lack of distinctive morphological traits [10,60].

Although a culture and genomic data for DEEPOLs are unavailable, some aspects of their biology can be inferred from the available evidence. Despite phylogeny and SSNs linking DEEPOLs to photosynthetic groups, we infer that DEEPOLs correspond to pico-to nanosized heterotrophic cells. This is based on their vertical distribution as they were consistently detected not only in sunlit layers but also in dark ocean samples, indicating that these lineages may not depend on light. For DEEPOL2 we have the additional support from *in situ* visualization which revealed no evidence of photosynthetic plastids, consistent with a heterotrophic, likely phagotrophic lifestyle. A comparable case is Picozoa, formerly known as picobiliphytes, which were first interpreted as photosynthetic algae related to cryptophytes and katablepharids [67], but were later shown by single-cell genomics and microscopy to lack plastids, even though they group within Archaeplastida [9,68,69]. Like the DEEPOLs, Picozoa occur in both sunlit and dark ocean waters [70]. The only described picozoan species was proposed to feed on nano-sized particles by endocytosis [69], so the DEEPOL feeding mode need not be classical grazing. However, the feeding mode of DEEPOLs cannot be resolved from our data.

While the DEEPOLs likely share a heterotrophic lifestyle, they appear to occupy distinct ecological niches. DEEPOL1 shows a cosmopolitan distribution across ocean basins and depth layers without a consistent preference for coastal or open-ocean waters. Such a pattern is consistent with an environmentally flexible lineage with broad ecological tolerance and suggests limited habitat-specific environmental filtering, with occurrence shaped primarily by large-scale processes such as ocean circulation and current-driven dispersal [60]. Despite this broad distribution, DEEPOL1 remains persistently rare, suggesting that factors other than habitat availability constrain its population size. Rather than reflecting habitat-driven selection, this rarity may instead result from selection acting directly on abundance. For example, selection favoring low population densities as a means of escaping predation or parasitism can maintain taxa at low abundance even where conditions are otherwise favorable [10]. DEEPOL3 also appears to represent a permanently rare lineage, but, unlike DEEPOL1, it shows a habitat-restricted distribution (Fig. 2C, 5C). Its consistently low abundance and restriction to a defined set of conditions suggest ecological specialization rather than the stochastic presence expected of a dispersal-derived transient taxon [71,72]. This points to a rarity driven by habitat specialization and narrow niche breadth, rather than by selection acting on abundance directly as hypothesized for DEEPOL1. Rare taxa of this type have been hypothesized to contribute specific ecological functions that are maintained at low population sizes across space and time [58,60]. DEEPOL2 exhibits yet another pattern. Across most of its range it is detected slightly more often offshore, but this observation reverses in polar waters (Fig. 3). DEEPOL2 also shows the capacity for pronounced local or regional increases in abundance, most notably at high southern latitudes (Fig. 2B), although this signal is driven predominantly by a single ASV (Fig. 5B). This ASV reaches high relative abundance independently in two geographically distinct regions, the Western Antarctic Peninsula shelf and the sub-Antarctic Pacific south of New Zealand, suggesting that this blooming capacity is an inherent trait of this DEEPOL2 ASV rather than a single localized event. These observations are consistent with a conditionally rare taxon that persists at low abundance across much of its range but becomes abundant under specific environmental conditions [73]. Such lineages are often interpreted as components of a microbial “seed bank,” remaining rare with low activity until favourable conditions, such as elevated nutrient availability or shifts in prey populations, allow for rapid population growth [60,74].

To better understand the evolutionary origins of the niche patterns in the DEEPOLs, we examined how niche categories (polar vs. non-polar; sunlit vs. dark ocean) and coastal vs. open-ocean indicator ASVs are distributed across each lineage’s ASV-level phylogeny (Fig. 5). Overall, qualitative inspection of the phylogeny and niche associations suggests a pattern more consistent with phylogenetic overdispersion than phylogenetic niche conservatism across the three DEEPOL lineages [75]. This is particularly evident for the polar niche in DEEPOL1, where the two polar-specialized ASVs occupy very distant positions within the tree (Fig. 5A). A similar pattern has previously been observed in Picozoa [70], and it has been hypothesized that specific ecological dynamics in polar ecosystems select for a diverse, distantly related range of organisms capable of coping with challenging conditions [76,77]. The polar niche in DEEPOL2 shows a different pattern, with polar-specialized and polar-generalist ASVs restricted to a single clade, which may indicate phylogenetic niche conservatism (Fig. 5B) [75]. However, the clade is poorly resolved and we lack niche information for some of the sequences. The polar-dominated subclade also contains a nested group of exclusively non-polar specialists, indicating that ecological divergence between polar and non-polar niches occurred within this lineage. Specialization to the dark ocean environment also largely follows a pattern of phylogenetic overdispersion in all three lineages, although a certain degree of conservatism can be observed in subclades composed of dark-ocean specialists specifically in DEEPOL3 (Fig. 5C). Because the sunlit specialists in DEEPOL3 are concentrated in a single clade, the current resolution of the tree suggests that the sunlit preference may have evolved from a dark-ocean specialized common ancestor, although this would need to be tested with a better-resolved phylogeny and formal ancestral state reconstruction. Several factors could underlie the phylogenetic overdispersion observed in the DEEPOLs, with competitive exclusion, ecological divergence, and convergent adaptation representing the main potential drivers of this pattern [78,79]. Which of these mechanisms drives niche differentiation among DEEPOL ASVs cannot be determined from a metabarcoding dataset alone. However, if the DEEPOLs are heterotrophic, as their morphology suggests, the same reasoning applied to Picozoa [70] may also hold here. In heterotrophs, phylogenetically conserved traits such as feeding-apparatus size lead related species to share prey preferences, intensifying competition and making mutual exclusion from the same niche more likely.

## Conclusion

We describe three orphan protist lineages recovered from long-read environmental sequencing and, for one of them, provide the first images of the cells themselves. The combined approach used here, coupling long-read metabarcoding and sequence similarity networks with global short-read surveys, lineage-specific dPCR and CARD-FISH, offers a route to identify and place rare and orphan protist lineages into a preliminary phylogenetic and ecological context. The putative placement of DEEPOLs in the vicinity of plastid-containing groups makes them potentially relevant to questions extending beyond their own biology, including plastid evolution. Recent examples of heterotrophic lineages branching close to photosynthetic groups, such as *Rhodelphis* and Picozoa, have shed new light into the convoluted evolution of plastids [3,9,80]. More broadly, the recovery of three previously unrecognized orphan lineages from a single long-read dataset suggests that the rare biosphere holds further uncharacterized diversity, and that the limiting factor is no longer sequencing effort but the ability to connect divergent sequences to cells.

## Acknowledgements

We thank Dolors Vaqué and Ramon Massana for providing us with the samples from the Southern Ocean. We acknowledge the donation made in Birgitta Sintring’s name to the Competence Center for Hidden Biodiversity, Department of Ecology and Genetics, Uppsala University, which enabled the confocal microscopy. In preparation of this manuscript, we used Claude Sonnet 5.5 (Anthropic) to revise and debug parts of the analysis code (R). We reviewed AI-generated content and take full responsibility for the content of this manuscript.

## Data availability

The long-read metabarcoding sequences and tree used to identify new orphan lineages can be found under https://doi.org/10.6084/m9.figshare.15164772.v3. The ASV table and metadata from the EukBank dataset used here for biogeographical and ecological analyses can be found under https://zenodo.org/records/7804946. The scripts used for the inference of SSNs is available under https://github.com/MiguelMSandin/SSNetworks. All code for the biogeography analysis of the DEEPOL lineages is published under https://github.com/burki-lab/deep-orphan-lineages.

## References

1. Burki F, Roger AJ, Brown MW et al. The New Tree of Eukaryotes. Trends in Ecology & Evolution 2020;35(1):43–55. 10.1016/j.tree.2019.08.008.

2. Eglit Y, Shiratori T, Jerlström-Hultqvist J et al. Meteora sporadica, a protist with incredible cell architecture, is related to Hemimastigophora. Current Biology 2024;34(2):451–459.e6. 10.1016/j.cub.2023.12.032.

3. Gawryluk RMR, Tikhonenkov DV, Hehenberger E et al. Non-photosynthetic predators are sister to red algae. Nature 2019;572(7768):art. 7768. 10.1038/s41586-019-1398-6.

4. Janouškovec J, Tikhonenkov DV, Burki F et al. A New Lineage of Eukaryotes Illuminates Early Mitochondrial Genome Reduction. Current Biology 2017;27(23):3717–3724.e5. 10.1016/j.cub.2017.10.051.

5. Worden AZ, Follows MJ, Giovannoni SJ et al. Rethinking the marine carbon cycle: Factoring in the multifarious lifestyles of microbes. Science 2015;347(6223):1257594. 10.1126/science.1257594.

6. López-García P, Moreira D. Open Questions on the Origin of Eukaryotes. Trends in Ecology & Evolution 2015;30(11):697–708. 10.1016/j.tree.2015.09.005.

7. Caron DA, Countway PD, Jones AC et al. Marine Protistan Diversity. Annual Review of Marine Science 2012;4(Volume 4, 2012):467–93. 10.1146/annurev-marine-120709-142802.

8. Valt M, Pánek T, Mirzoyan S et al. Rare microbial relict sheds light on an ancient eukaryotic supergroup. Nature 2026;649(8096):388–95. 10.1038/s41586-025-09750-0.

9. Schön ME, Zlatogursky VV, Singh RP et al. Single cell genomics reveals plastid-lacking Picozoa are close relatives of red algae. Nat Commun 2021;12(1):art. 1. 10.1038/s41467-021-26918-0.

10. Logares R, Mangot JF, Massana R. Rarity in aquatic microbes: placing protists on the map. Research in Microbiology Special issue on Microbial diversity, adaptation and evolution, 2015;166(10):831–41. 10.1016/j.resmic.2015.09.009.

11. López-Escardó D, Obiol A, Marimon G et al. Single-cell sequencing reveals genome streamlining and functional diversity in ecologically dominant marine protists. Preprint, bioRxiv, 21 Apr. 2026. 10.64898/2026.04.17.719054.

12. Wideman JG, Monier A, Rodríguez-Martínez R et al. Unexpected mitochondrial genome diversity revealed by targeted single-cell genomics of heterotrophic flagellated protists. Nat Microbiol 2020;5(1):154–65. 10.1038/s41564-019-0605-4.

13. Brown CT, Hug LA, Thomas BC et al. Unusual biology across a group comprising more than 15% of domain Bacteria. Nature 2015;523(7559):208–11. 10.1038/nature14486.

14. Parks DH, Rinke C, Chuvochina M et al. Recovery of nearly 8,000 metagenome-assembled genomes substantially expands the tree of life. Nat Microbiol 2017;2(11):1533–42. 10.1038/s41564-017-0012-7.

15. Gong X, Río ÁR del, Xu L et al. New globally distributed bacterial phyla within the FCB superphylum. Nat Commun 2022;13(1):7516. 10.1038/s41467-022-34388-1.

16. Spang A, Saw JH, Jørgensen SL et al. Complex archaea that bridge the gap between prokaryotes and eukaryotes. Nature 2015;521(7551):173–9. 10.1038/nature14447.

17. Zaremba-Niedzwiedzka K, Caceres EF, Saw JH et al. Asgard archaea illuminate the origin of eukaryotic cellular complexity. Nature 2017;541(7637):353–8. 10.1038/nature21031.

18. Jamy M, Huber T, Antoine T et al. Identification of a deep-branching lineage of algae using environmental plastid genomes. Nat Commun 2025;17(1):662. 10.1038/s41467-025-67401-4.

19. Serra Moncadas L, Ruscheweyh HJ, Woodhouse J et al. A global mitochondrial genomic atlas illuminates freshwater microeukaryote diversity. Nat Commun published online 25 Sept. 2026. 10.1038/s41467-026-78160-1.

20. Delmont TO, Gaia M, Hinsinger DD et al. Functional repertoire convergence of distantly related eukaryotic plankton lineages abundant in the sunlit ocean. Cell Genomics 2022;2(5):100123. 10.1016/j.xgen.2022.100123.

21. Cordier T, Angeles IB, Henry N et al. Patterns of eukaryotic diversity from the surface to the deep-ocean sediment. Science Advances 2022;8(5):eabj9309. 10.1126/sciadv.abj9309.

22. Vargas C de, Audic S, Henry N et al. Eukaryotic plankton diversity in the sunlit ocean. Science 2015;348(6237):1261605. 10.1126/science.1261605.

23. Dunthorn M, Otto J, Berger SA et al. Placing Environmental Next-Generation Sequencing Amplicons from Microbial Eukaryotes into a Phylogenetic Context. Molecular Biology and Evolution 2014;31(4):993–1009. 10.1093/molbev/msu055.

24. Jamy M, Foster R, Barbera P et al. Long-read metabarcoding of the eukaryotic rDNA operon to phylogenetically and taxonomically resolve environmental diversity. Molecular Ecology Resources 2020;20(2):429–43. 10.1111/1755-0998.13117.

25. Edgar RC, Haas BJ, Clemente JC et al. UCHIME improves sensitivity and speed of chimera detection. Bioinformatics 2011;27(16):2194–200. 10.1093/bioinformatics/btr381.

26. Callahan BJ, McMurdie PJ, Rosen MJ et al. DADA2: High-resolution sample inference from Illumina amplicon data. Nat Methods 2016;13(7):581–3. 10.1038/nmeth.3869.

27. Burki F, Sandin MM, Jamy M. Diversity and ecology of protists revealed by metabarcoding. Current Biology 2021;31(19):R1267–80. 10.1016/j.cub.2021.07.066.

28. Jamy M, Biwer C, Vaulot D et al. Global patterns and rates of habitat transitions across the eukaryotic tree of life. Nat Ecol Evol 2022;6(10):art. 10. 10.1038/s41559-022-01838-4.

29. Thoen E, Krabberød AK, Nogerius VT et al. Short- and long-read metabarcoding of Archaeorhizomycetes reveals high phylogenetic diversity structured by vegetation and climate. New Phytologist 2026;249(1):418–32. 10.1111/nph.70682.

30. Chwalińska M, Karlicki M, Romac S et al. From short to long reads: enhanced protist diversity profiling via Nanopore metabarcoding. Metabarcoding and Metagenomics 2025;9:e163750. 10.3897/mbmg.9.163750.

31. Katoh K, Standley DM. MAFFT Multiple Sequence Alignment Software Version 7: Improvements in Performance and Usability. Molecular Biology and Evolution 2013;30(4):772–80. 10.1093/molbev/mst010.

32. Capella-Gutiérrez S, Silla-Martínez JM, Gabaldón T. trimAl: a tool for automated alignment trimming in large-scale phylogenetic analyses. Bioinformatics 2009;25(15):1972–3. 10.1093/bioinformatics/btp348.

33. Minh BQ, Schmidt HA, Chernomor O et al. IQ-TREE 2: New Models and Efficient Methods for Phylogenetic Inference in the Genomic Era. Molecular Biology and Evolution 2020;37(5):1530–4. 10.1093/molbev/msaa015.

34. Berney C, Mahé F, Henry N et al. EukBank 18S V4 dataset. Zenodo, 30 Oct. 2023. 10.5281/zenodo.7804946.

35. Flanders Marine Institute. IHO Sea Areas, version 3. Flanders Marine Institute (VLIZ), 2018. 10.14284/323.

36. Pebesma E. The R Journal: Simple Features for R: Standardized Support for Spatial Vector Data. *The R Journal* 2018;10(1):439–46. 10.32614/RJ-2018-009.

37. Cáceres MD, Legendre P. Associations between species and groups of sites: indices and statistical inference. Ecology 2009;90(12):3566–74. 10.1890/08-1823.1.

38. R Core Team. *R: A Language and Environment for Statistical Computing*, version 4.5.1. R Foundation for Statistical Computing, 2025. https://www.r-project.org/.

39. Wickham H, Averick M, Bryan J et al. Welcome to the Tidyverse. Journal of Open Source Software 2019;4(43):1686. 10.21105/joss.01686.

40. Wickham H. Ggplot2. Use R! Cham: Springer International Publishing, 2016. 10.1007/978-3-319-24277-4.

41. Oksanen J, Kindt R, Legendre P et al. The Vegan Package. 19 Jan. 2009.

42. Paradis E, Schliep K. *Ape 5.0: An Environment for Modern Phylogenetics and Evolutionary Analyses in R*. n.d. 10.1093/bioinformatics/bty633 (4 May 2026, date last accessed).

43. Yu G, Smith DK, Zhu H et al. ggtree: an r package for visualization and annotation of phylogenetic trees with their covariates and other associated data. Methods in Ecology and Evolution 2017;8(1):28–36. 10.1111/2041-210X.12628.

44. Conway JR, Lex A, Gehlenborg N. UpSetR: an R package for the visualization of intersecting sets and their properties. Bioinformatics 2017;33(18):2938–40. 10.1093/bioinformatics/btx364.

45. Massicotte, P.; South, A. Rnaturalearth: World Map Data from Natural Earth, version 1.2.0. 2026. 10.32614/CRAN.package.rnaturalearth.

46. Robinson, D.; Hayes, A.; Couch, S.; Hvitfeldt, E. Broom: Convert Statistical Objects into Tidy Tibbles, version 1.0.12. 2026. 10.32614/CRAN.package.broom.

47. Sandin MM, Walde M, Henry N et al. OligoN-Design: A Simple and Versatile Tool to Design Specific Probes and Primers From Large Heterogeneous Datasets. Molecular Ecology Resources 2026;26(3):e70140. 10.1111/1755-0998.70140.

48. Vaqué D, Guadayol Ò, Peters F et al. Differential response of grazing and bacterial heterotrophic production to experimental warming in Antarctic waters. Aquat Microb Ecol 2009;54:101–12. 10.3354/ame01259.

49. Pernice MC, Forn I, Gomes A et al. Global abundance of planktonic heterotrophic protists in the deep ocean. The ISME Journal 2015;9(3):782–92. 10.1038/ismej.2014.168.

50. Tikhonenkov DV, Mikhailov KV, Gawryluk RMR et al. Microbial predators form a new supergroup of eukaryotes. Nature 2022;612(7941):art. 7941. 10.1038/s41586-022-05511-5.

51. Burki F. The Eukaryotic Tree of Life from a Global Phylogenomic Perspective. Cold Spring Harb Perspect Biol 2014;6(5):a016147. 10.1101/cshperspect.a016147.

52. Campo J del, Kolisko M, Boscaro V et al. EukRef: Phylogenetic curation of ribosomal RNA to enhance understanding of eukaryotic diversity and distribution. PLOS Biology 2018;16(9):e2005849. 10.1371/journal.pbio.2005849.

53. Philippe* and H, Germot A. Phylogeny of Eukaryotes Based on Ribosomal RNA: Long-Branch Attraction and Models of Sequence Evolution. Mol Biol Evol 2000;17(5):830–4. 10.1093/oxfordjournals.molbev.a026362.

54. Arroyo AS, Lannes R, Bapteste E et al. Gene Similarity Networks Unveil a Potential Novel Unicellular Group Closely Related to Animals from the Tara Oceans Expedition. Genome Biology and Evolution 2020;12(9):1664–78. 10.1093/gbe/evaa117.

55. Forster D, Bittner L, Karkar S et al. Testing ecological theories with sequence similarity networks: marine ciliates exhibit similar geographic dispersal patterns as multicellular organisms. BMC Biology 2015;13(1):16. 10.1186/s12915-015-0125-5.

56. Angeles IB, Nguyen NL, Greco M et al. Assigning the unassigned: A signature-based classification of rDNA metabarcodes reveals new deep-sea diversity. PLOS ONE 2024;19(2):e0298440. 10.1371/journal.pone.0298440.

57. Tedersoo L, Mikryukov V, Sildever S et al. Discovering 25 novel phyla that fill gaps in the eukaryotic tree of life. Preprint, bioRxiv, 31 Aug. 2026, 2026.08.28.747736. 10.64898/2026.08.28.747736.

58. Jousset A, Bienhold C, Chatzinotas A et al. Where less may be more: how the rare biosphere pulls ecosystems strings. The ISME Journal 2017;11(4):853–62. 10.1038/ismej.2016.174.

59. Caron DA, Countway PD. Hypotheses on the role of the protistan rare biosphere in a changing world. Aquatic Microbial Ecology 2009;57(3):227–38. 10.3354/ame01352.

60. Lynch MDJ, Neufeld JD. Ecology and exploration of the rare biosphere. Nat Rev Microbiol 2015;13(4):217–29. 10.1038/nrmicro3400.

61. Decelle J, Carradec Q, Pochon X et al. Worldwide Occurrence and Activity of the Reef-Building Coral Symbiont Symbiodinium in the Open Ocean. Current Biology 2018;28(22):3625–3633.e3. 10.1016/j.cub.2018.09.024.

62. Yabuki A, Inagaki Y, Ishida K ichiro. *Palpitomonas bilix* gen. et sp. nov.: A Novel Deep-branching Heterotroph Possibly Related to Archaeplastida or Hacrobia. Protist 2010;161(4):523–38. 10.1016/j.protis.2010.03.001.

63. Leander BS. Eukaryotic evolution: Deep phylogeny does not imply morphological novelty. Current Biology 2023;33(3):R112–4. 10.1016/j.cub.2022.12.016.

64. Massana R, Balagué V, Guillou L et al. Picoeukaryotic diversity in an oligotrophic coastal site studied by molecular and culturing approaches. FEMS Microbiology Ecology 2004;50(3):231–43. 10.1016/j.femsec.2004.07.001.

65. Massana R, Pedrós-Alió C. Unveiling new microbial eukaryotes in the surface ocean. Current Opinion in Microbiology Ecology and Industrial Microbiology ● Techniques, 2008;11(3):213–8. 10.1016/j.mib.2008.04.004.

66. Obiol A, del Campo J, de Vargas C et al. How marine are Marine Stramenopiles (MAST)? A cross-system evaluation. FEMS Microbiology Ecology 2024;100(11):fiae130. 10.1093/femsec/fiae130.

67. Not F, Valentin K, Romari K et al. Picobiliphytes: A Marine Picoplanktonic Algal Group with Unknown Affinities to Other Eukaryotes. Science 2007;315(5809):253–5. 10.1126/science.1136264.

68. Yoon HS, Price DC, Stepanauskas R et al. Single-cell genomics reveals organismal interactions in uncultivated marine protists. Science 2011;332(6030):714–7. 10.1126/science.1203163.

69. Seenivasan R, Sausen N, Medlin LK et al. Picomonas judraskeda Gen. Et Sp. Nov.: The First Identified Member of the Picozoa Phylum Nov., a Widespread Group of Picoeukaryotes, Formerly Known as ‘Picobiliphytes’. PLOS ONE 2013;8(3):e59565. 10.1371/journal.pone.0059565.

70. Huber P, De Angelis D, Sarmento H et al. Global distribution, diversity, and ecological niche of Picozoa, a widespread and enigmatic marine protist lineage. Microbiome 2024;12(1):162. 10.1186/s40168-024-01874-1.

71. Pascoal F, Costa R, Magalhães C. The microbial rare biosphere: current concepts, methods and ecological principles. FEMS Microbiology Ecology 2021;97(1):fiaa227. 10.1093/femsec/fiaa227.

72. Jia X, Dini-Andreote F, Falcão Salles J. Community Assembly Processes of the Microbial Rare Biosphere. Trends in Microbiology 2018;26(9):738–47. 10.1016/j.tim.2018.02.011.

73. Shade A, Jones SE, Caporaso JG et al. Conditionally Rare Taxa Disproportionately Contribute to Temporal Changes in Microbial Diversity. mBio 2014;5(4):10.1128/mbio.01371-14. 10.1128/mbio.01371-14.

74. Lennon JT, Jones SE. Microbial seed banks: the ecological and evolutionary implications of dormancy. Nat Rev Microbiol 2011;9(2):119–30. 10.1038/nrmicro2504.

75. Losos JB. Phylogenetic niche conservatism, phylogenetic signal and the relationship between phylogenetic relatedness and ecological similarity among species. Ecology Letters 2008;11(10):995–1003. 10.1111/j.1461-0248.2008.01229.x.

76. Kleinteich J, Hildebrand F, Bahram M et al. Pole-to-Pole Connections: Similarities between Arctic and Antarctic Microbiomes and Their Vulnerability to Environmental Change. Front Ecol Evol 2017;5. 10.3389/fevo.2017.00137.

77. Rampelotto PH. Polar Microbiology: Recent Advances and Future Perspectives. Biology 2014;3(1):81–4. 10.3390/biology3010081.

78. Macarthur R, Levins R. The Limiting Similarity, Convergence, and Divergence of Coexisting Species. The American Naturalist 1967;101(921):377–85.

79. Violle C, Nemergut DR, Pu Z et al. Phylogenetic limiting similarity and competitive exclusion. Ecology Letters 2011;14(8):782–7. 10.1111/j.1461-0248.2011.01644.x.

80. Sibbald SJ, Archibald JM. Genomic Insights into Plastid Evolution. Genome Biology and Evolution 2020;12(7):978–90. 10.1093/gbe/evaa096.

