## Supplementary Figures for "Three novel deep-branching orphan protist lineages from the marine rare biosphere"

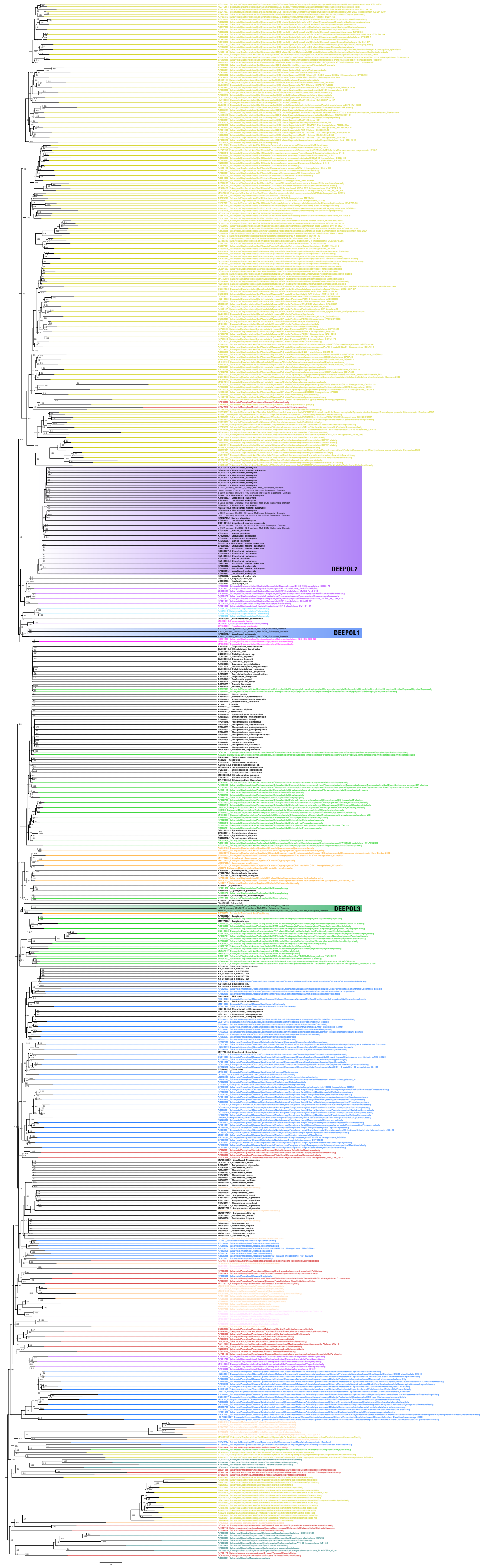
DEEPOL2
DEEPOL1
DEEPOL3

**Figure S1** Phylogenetic placement of the top NCBI nt/nr hits for each DEEPOL lineage based on the 18S rRNA gene. The backbone tree was built from sequences in the PR2 database. The maximum-likelihood tree was inferred with IQ-TREE under the GTR+F+R10 model, and node support values are ultrafast bootstrap values (1000 replicates). DEEPOL1, DEEPOL2 and DEEPOL3 are indicated by blue, purple and green rectangles, respectively. NCBI nt/nr hits are shown in bold.

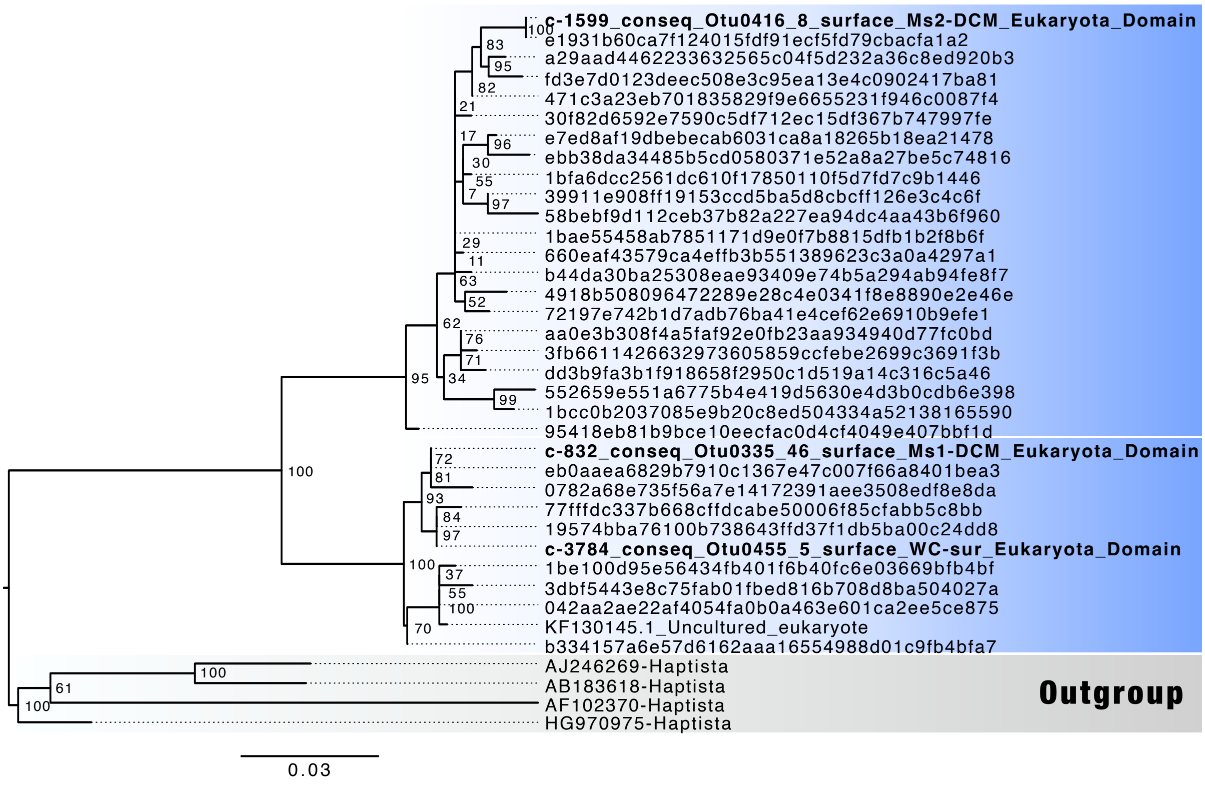


**Figure S2** 18S rDNA diversity of DEEPOL1. Maximum-likelihood phylogeny inferred from 18S rDNA sequences corresponding to DEEPOL1 under the TNe+R2 model, including long-read metabarcoding reference sequences and short-read ASVs recovered from the EukBank dataset. Putative subclades within DEEPOL1 are highlighted with blue rectangles, and the selected outgroup is indicated with a grey rectangle. Long-read metabarcoding sequences are shown in bold.

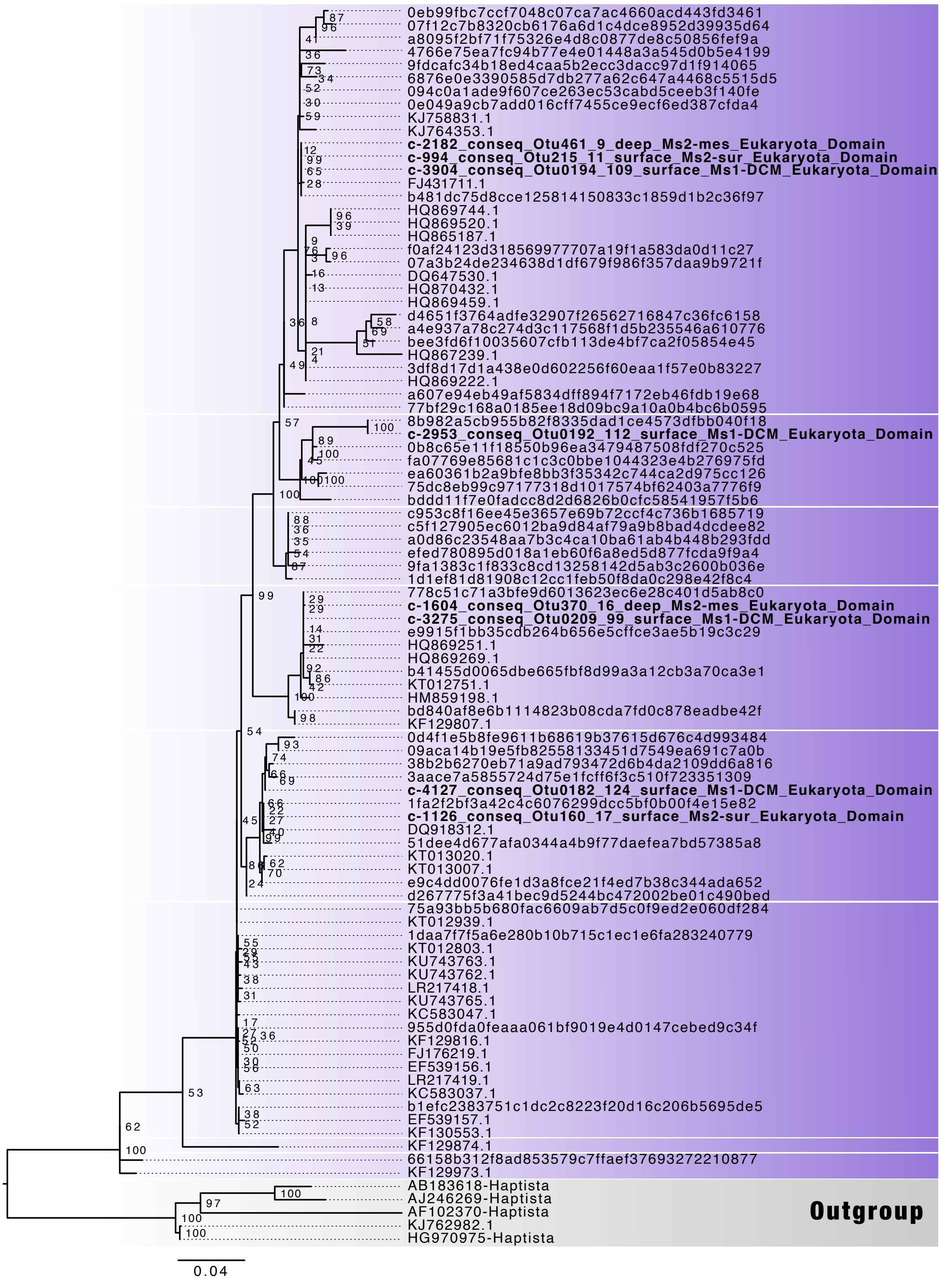


**Figure S3** 18S rDNA diversity of DEEPOL2. Maximum-likelihood phylogeny inferred from sequences corresponding to DEEPOL2 under the TNe+I+R2 model, including long-read metabarcoding sequences, additional 18S rDNA sequences from PR2 and NCBI as well as short-read ASVs recovered from the EukBank dataset. Putative subclades within DEEPOL2 are highlighted with purple rectangles, and the selected outgroup is indicated with a grey rectangle. Long-read metabarcoding sequences are shown in bold.

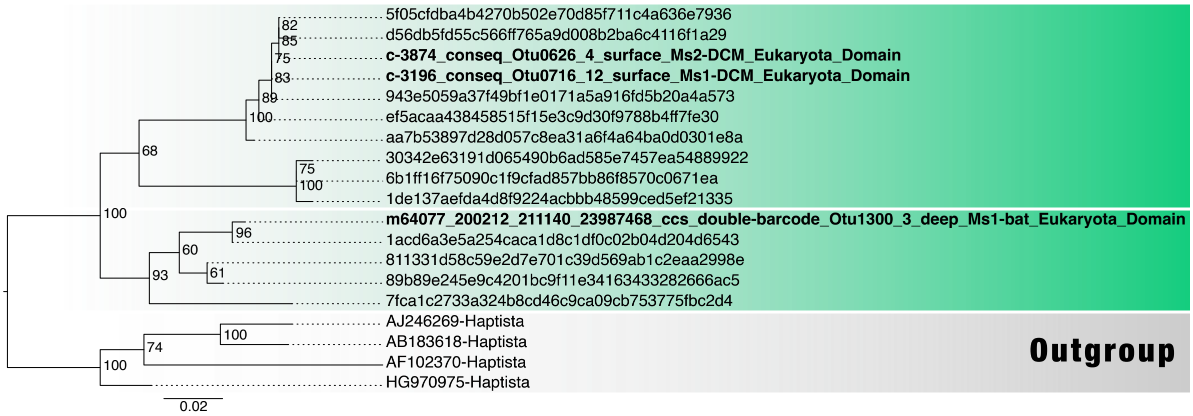


**Figure S4** 18S rDNA diversity of DEEPOL3. Maximum-likelihood phylogeny inferred from 18S rDNA sequences corresponding to DEEPOL3 under the TNe+R2 model, including long-read metabarcoding reference sequences and short-read ASVs recovered from the EukBank dataset. Putative subclades within DEEPOL3 are highlighted with green rectangles, and the selected outgroup is indicated with a grey rectangle. Long-read metabarcoding sequences are shown in bold.

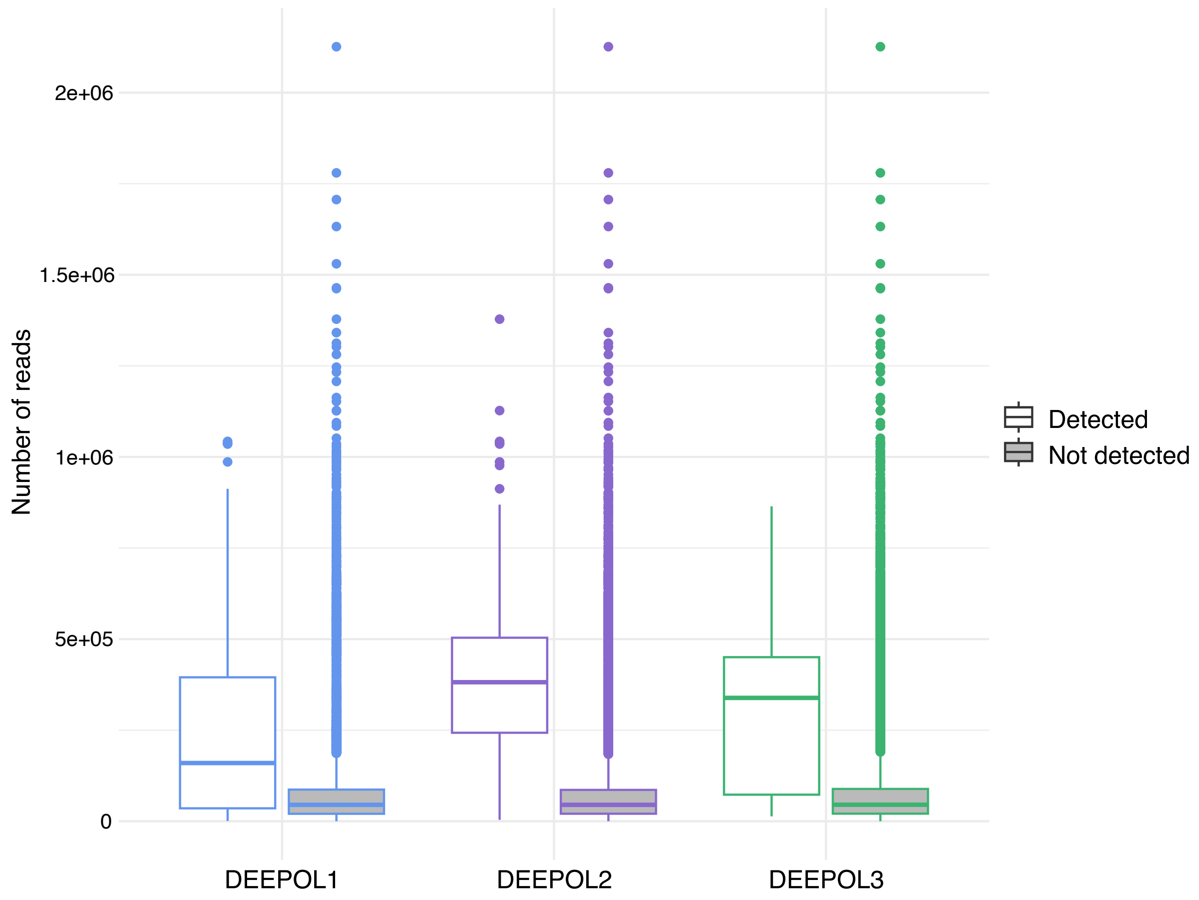


**Figure S5** Sequencing depth (number of reads) of EukBank samples in which each DEEPOL was detected and not detected. Boxes show the interquartile range and the median, whiskers extend to 1.5 times the interquartile range, and points are outliers

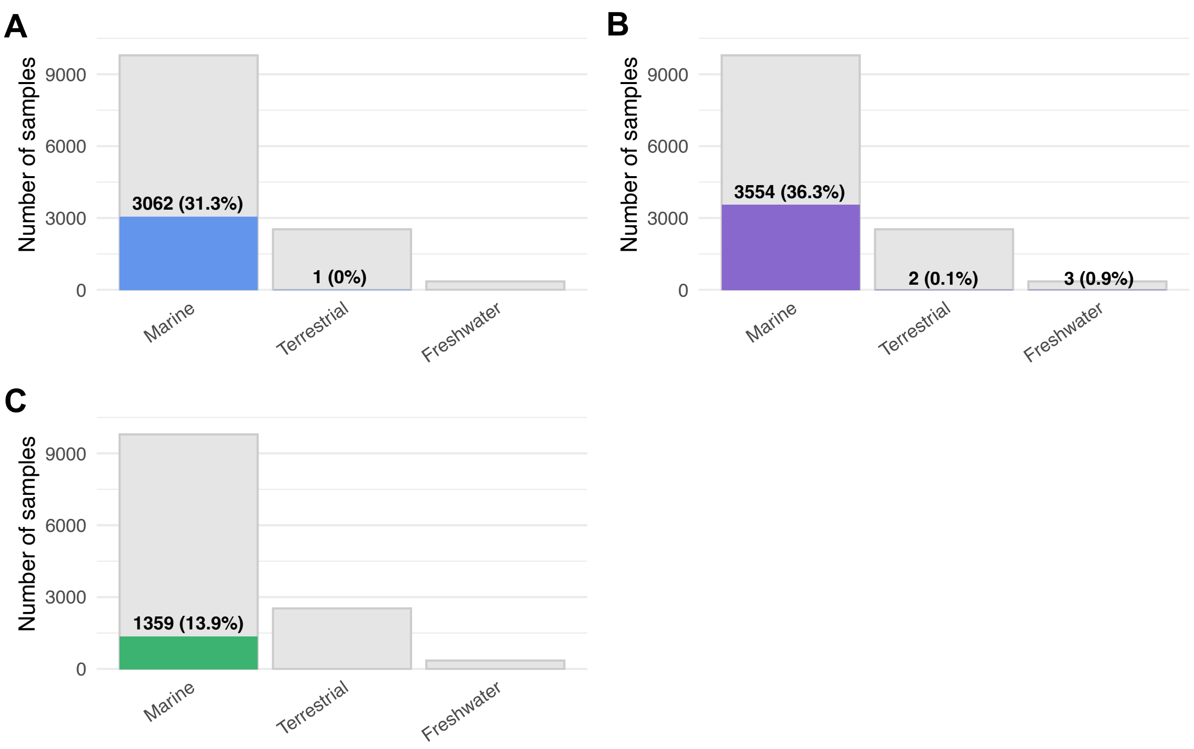


**Figure S6** Habitat occupancy of the DEEPOL lineages. Bar plots summarizing the number of samples in which DEEPOL1 (**A**), DEEPOL2 (**B**), and DEEPOL3 (**C**) were detected across different habitat categories.

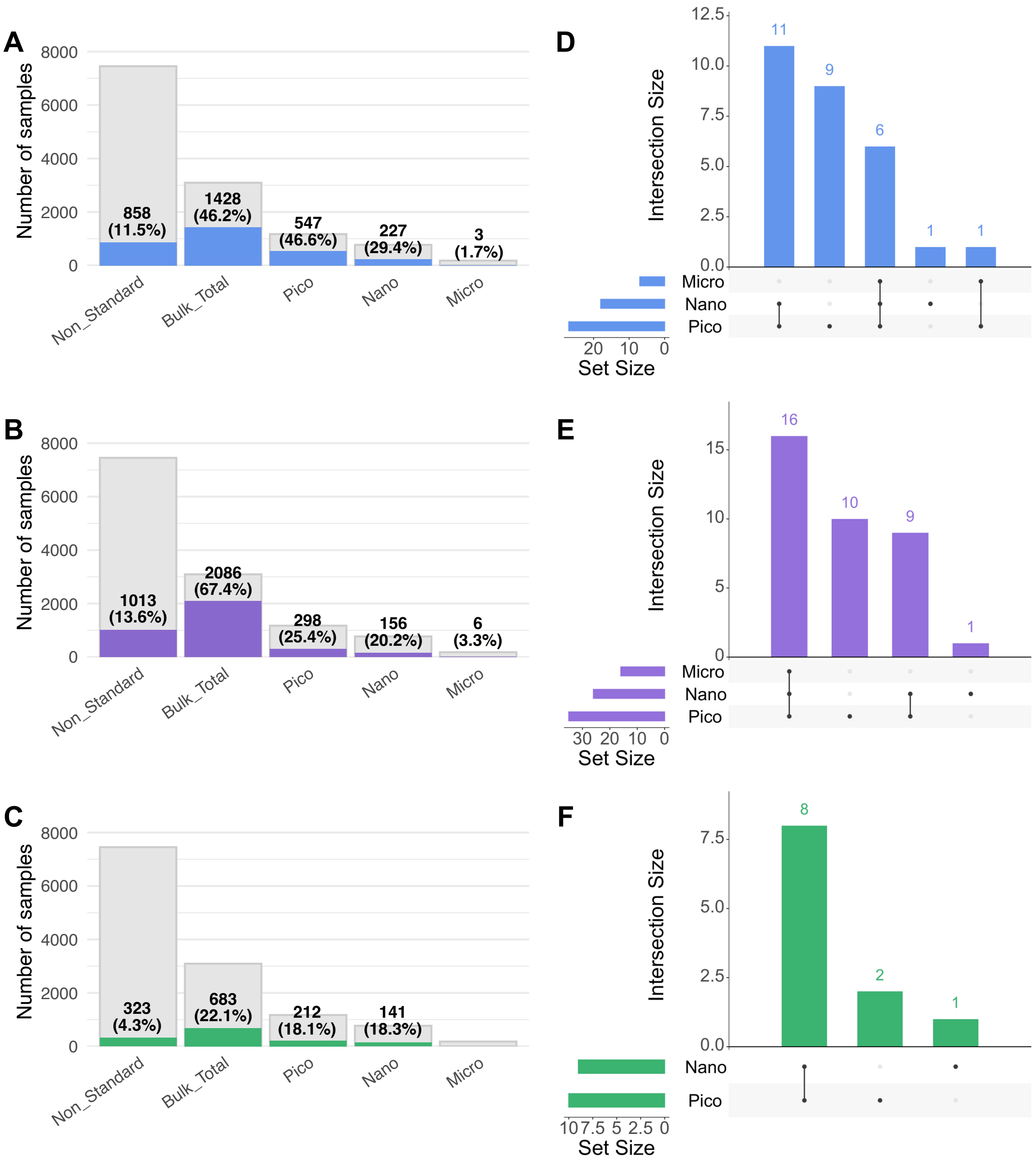


**Figure S7** Size-fraction occupancy of DEEPOL lineages and overlap of ASVs across size fractions. Bar plots summarizing the number of samples in which DEEPOL1 (**A**), DEEPOL2 (**B**), and DEEPOL3 (**C**) were detected across different and size fractions. UpSet plots illustrate overlap of ASVs among pico-, nano-, and micro-plankton size fractions for DEEPOL1 (**D**), DEEPOL2 (**E**), and DEEPOL3 (**F**).

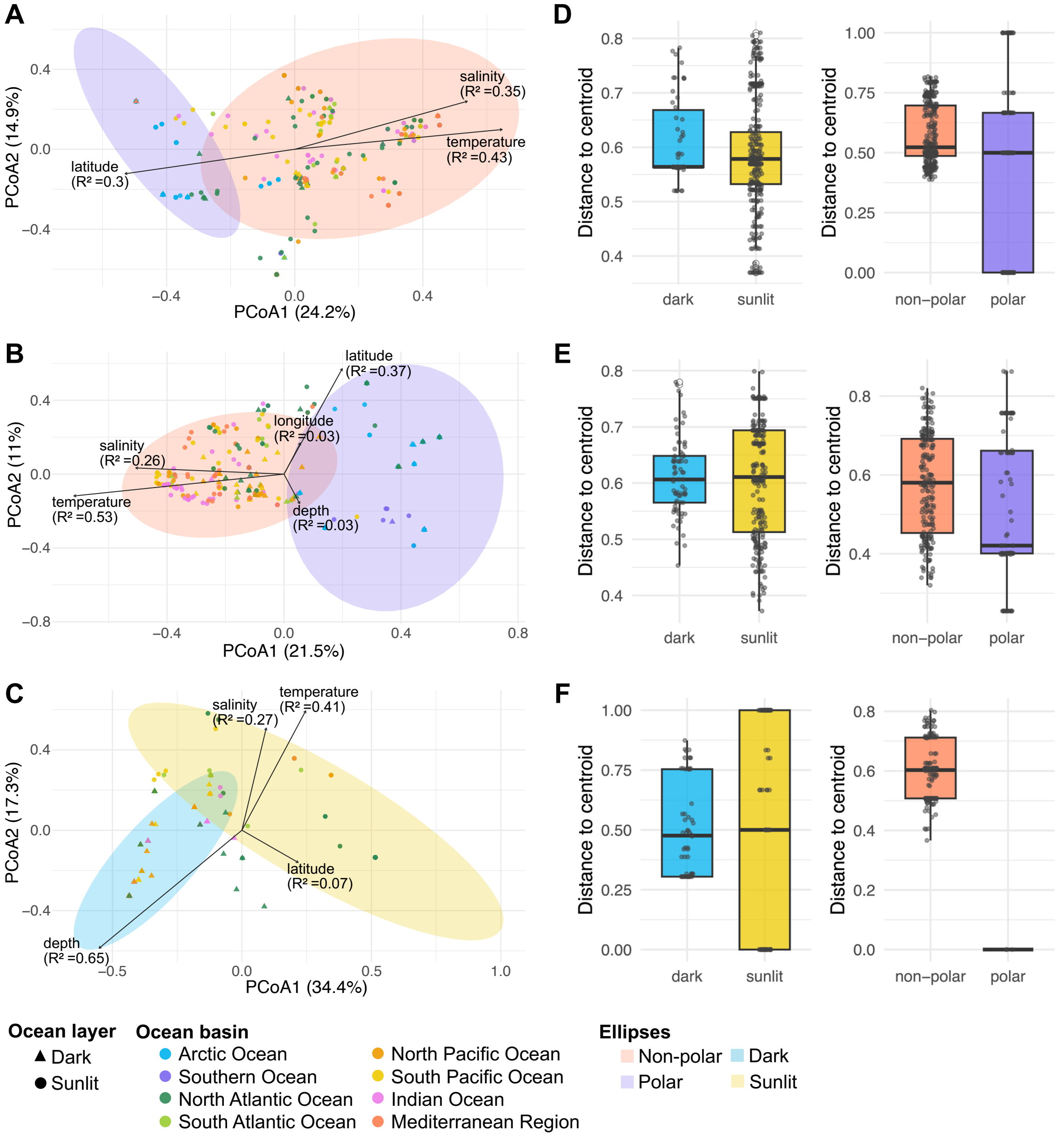


**Figure S8** Supplementary multivariate analyses of DEEPOL lineages. PCoA based on Jaccard dissimilarity illustrates major patterns in lineage-specific community composition for DEEPOL1, DEEPOL2, and DEEPOL3 (**A-C**). Boxplots showing within-group dispersion of samples used in the PERMDISP analyses are shown for polar versus non-polar ocean basins and for sunlit versus dark ocean layers for DEEPOL1–3 (**D-F**).

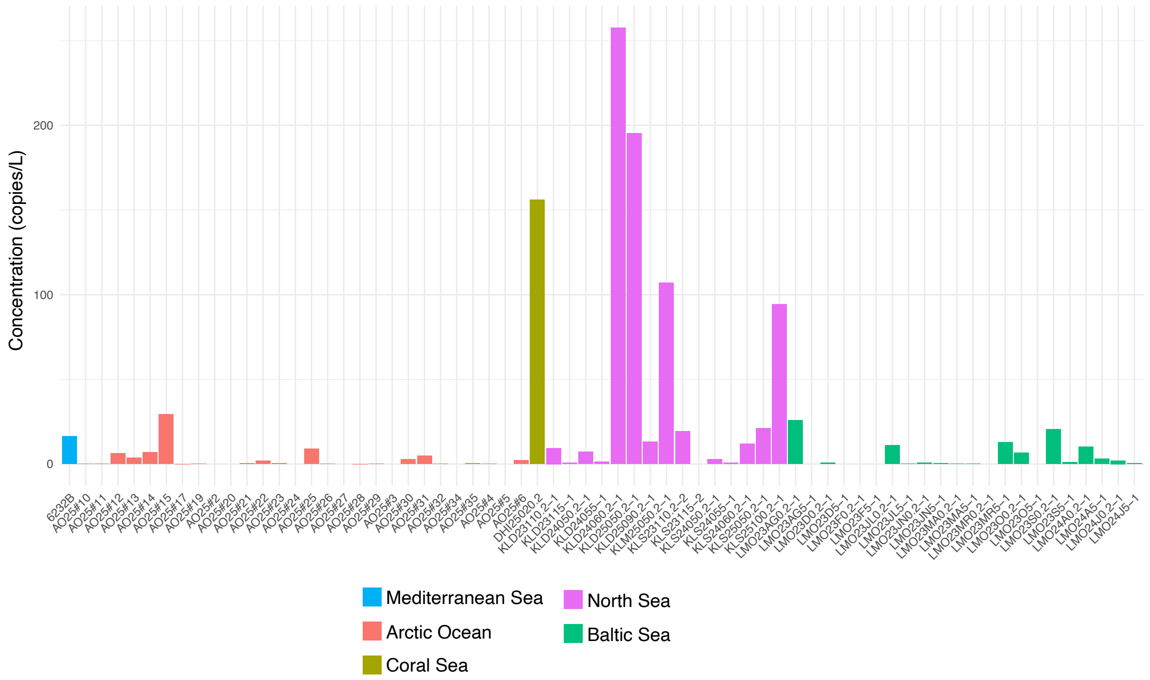


**Figure S9** Environmental abundance of DEEPOL2 inferred from digital PCR screening. Bar plot showing concentrations of DEEPOL2 18S rDNA (copies L$^{-1}$) measured by dPCR across 67 environmental DNA samples. Samples are ordered by sampling site.

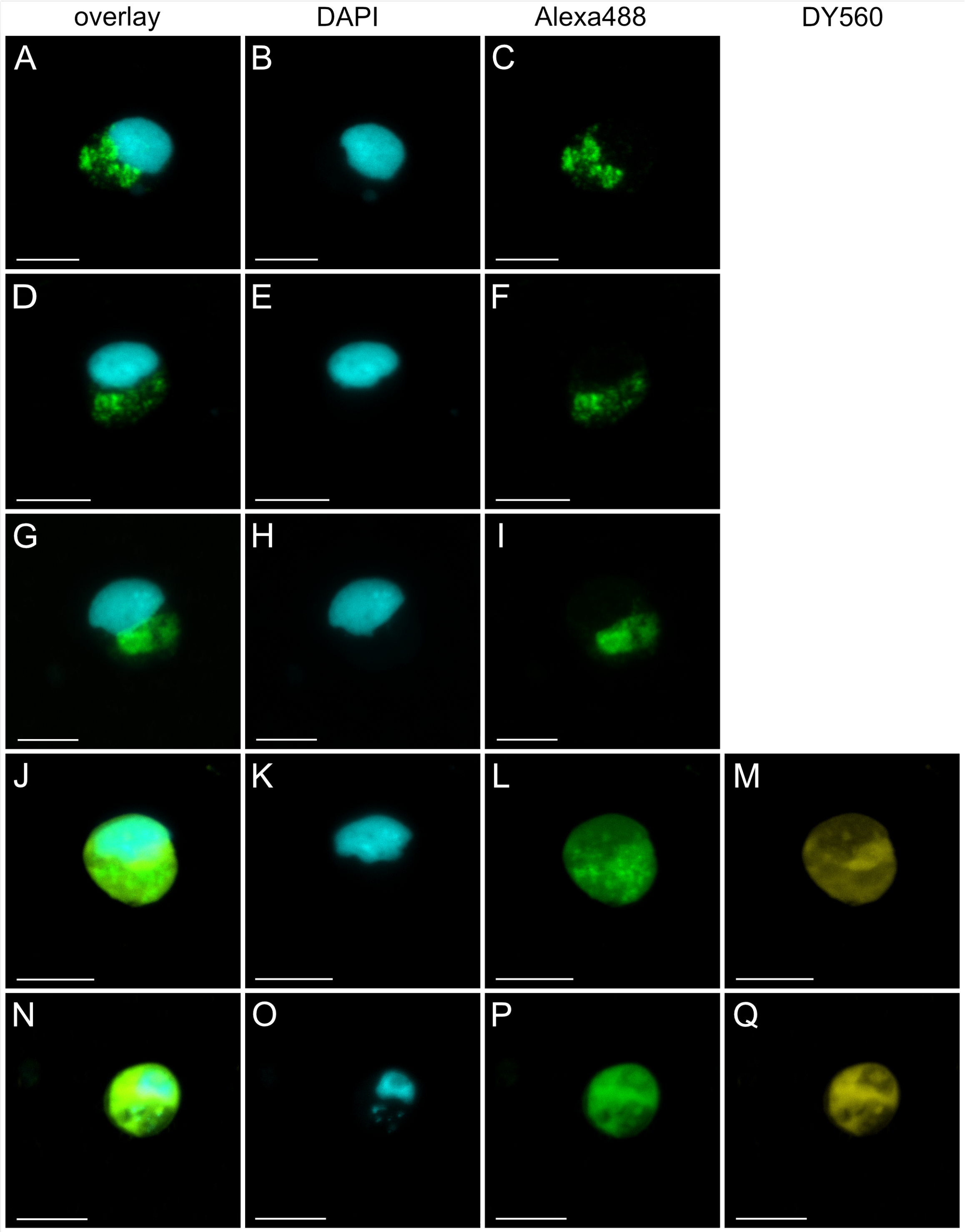


**Figure S10** *In situ* visualization of DEEPOL2 cells from the North Sea (Skagerrak). All CARD-FISH images acquired for Skagerrak samples are shown. Panels display overlays of all fluorescence channels (**A, D, G, J, N**), DAPI-stained nucleic acids (**B, E, H, K, O**), DEEPOL2-specific CARD-FISH probe signal detected with Alexa Fluor 488 (**C, F, I, L, P**), and NHS-ester protein staining using DY-560 (**M, Q**). Images shown represent maximum-intensity projections of Z-stacks acquired for each cell.

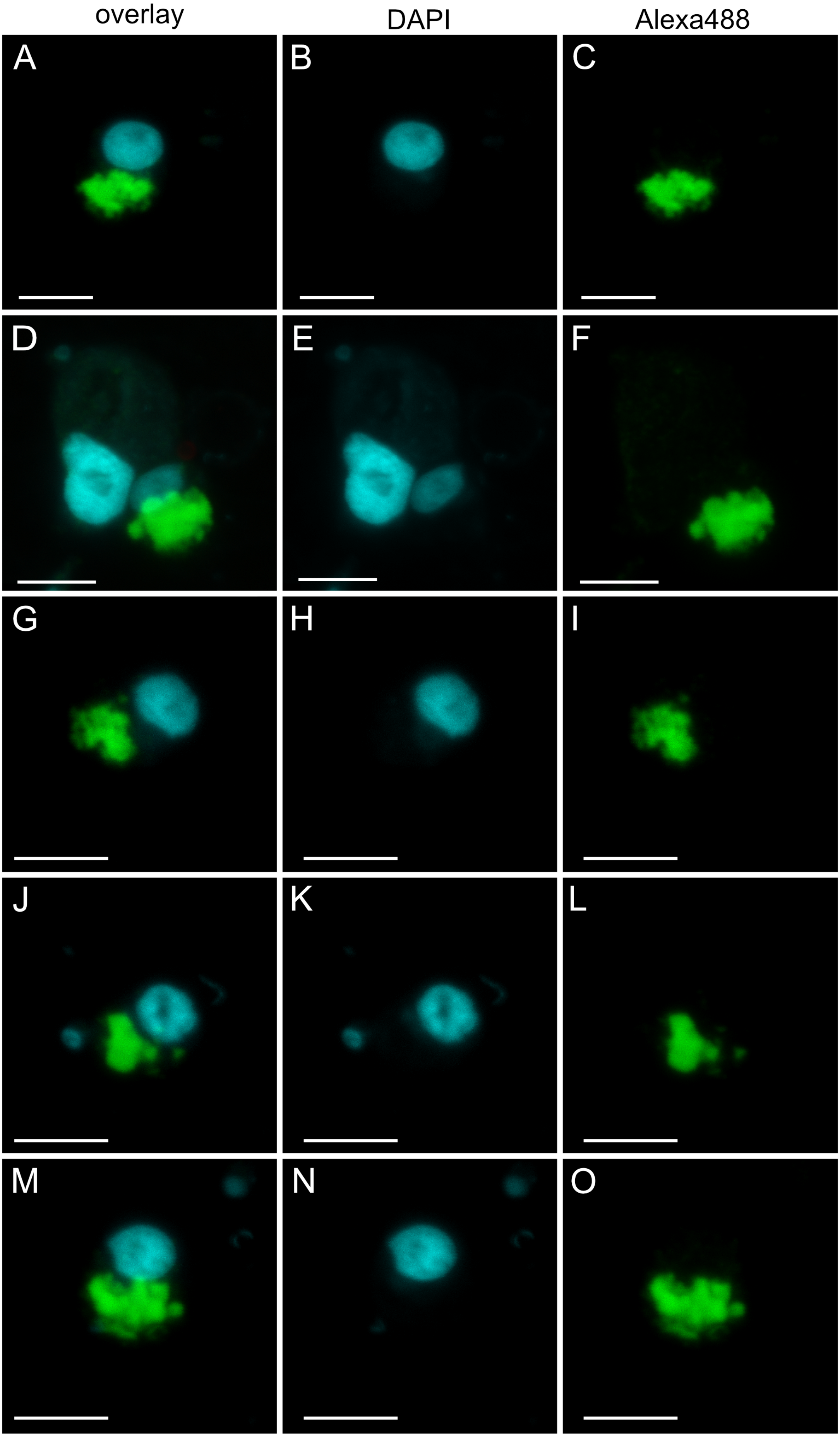

overlay
DAPI
Alexa488
A
B
C
D
E
F
G
H
I
J
K
L
M
N
O

**Figure S11** *In situ* visualization of DEEPOL2 cells from the Southern Ocean (Bellingshausen Sea). All CARD-FISH images acquired for Southern Ocean samples without NHS-ester are shown. Panels display overlays of all fluorescence channels (**A, D, G, J, M**), DAPI-stained nucleic acids (**B, E, H, K, N**), DEEPOL2-specific CARD-FISH probe signal detected with Alexa Fluor 488 (**C, F, I, L, O**). Images shown represent maximum-intensity projections of Z-stacks

acquired for each cell.

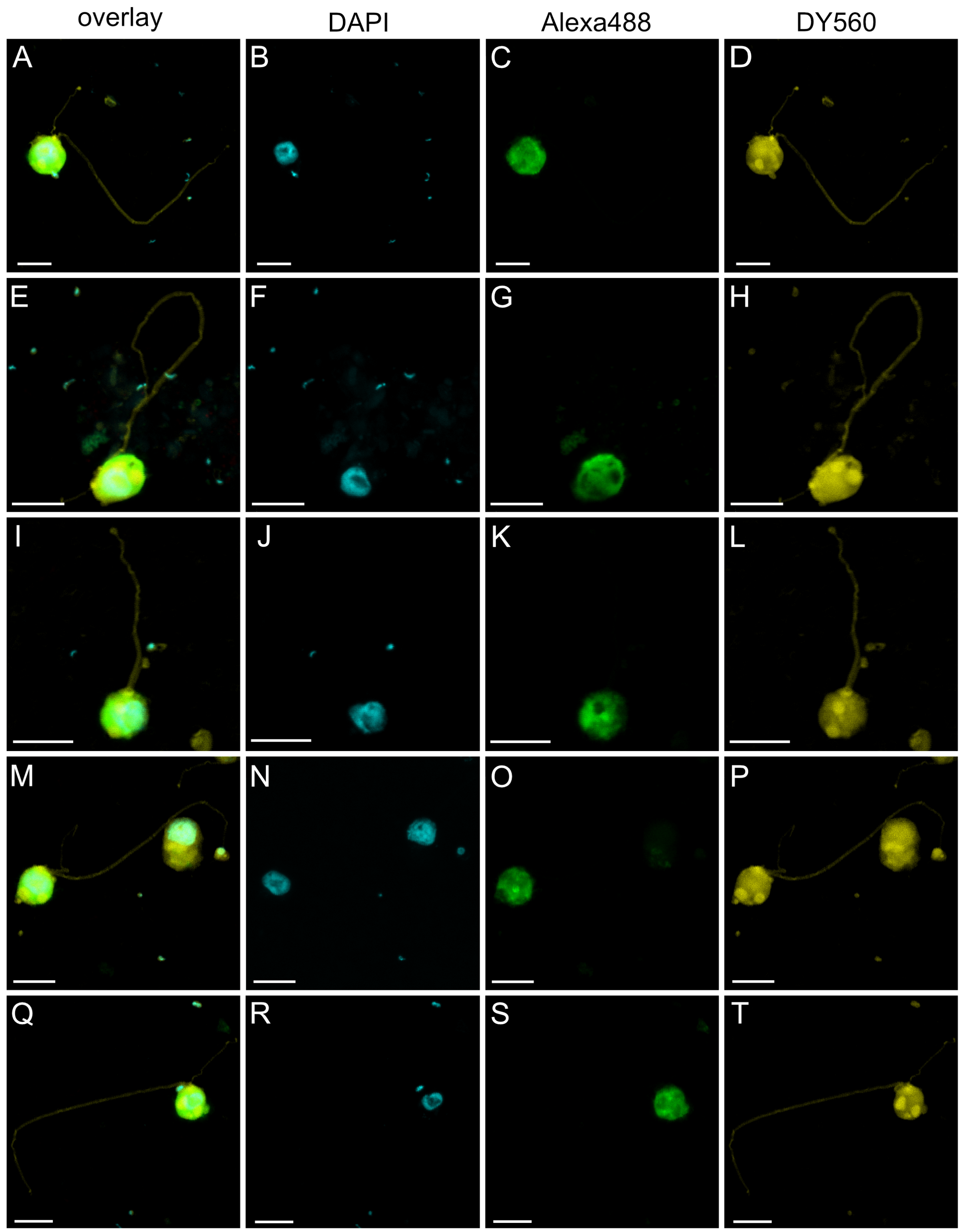


**Figure S12** *In situ* visualization of DEEPOL2 cells from the Southern Ocean.
All CARD-FISH images acquired for Southern Ocean samples with NHS-ester are shown. Panels display overlays of all fluorescence channels (**A, E, I, M, Q**), DAPI-stained nucleic acids (**B, F, J, N, R**), DEEPOL2-specific CARD-FISH probe signal detected with Alexa 488

(**C, G, K, O, S**), and NHS-ester protein staining using DY-560 (**D, H, L, P, T**). Images shown represent maximum-intensity projections of Z-stacks acquired for each cell.

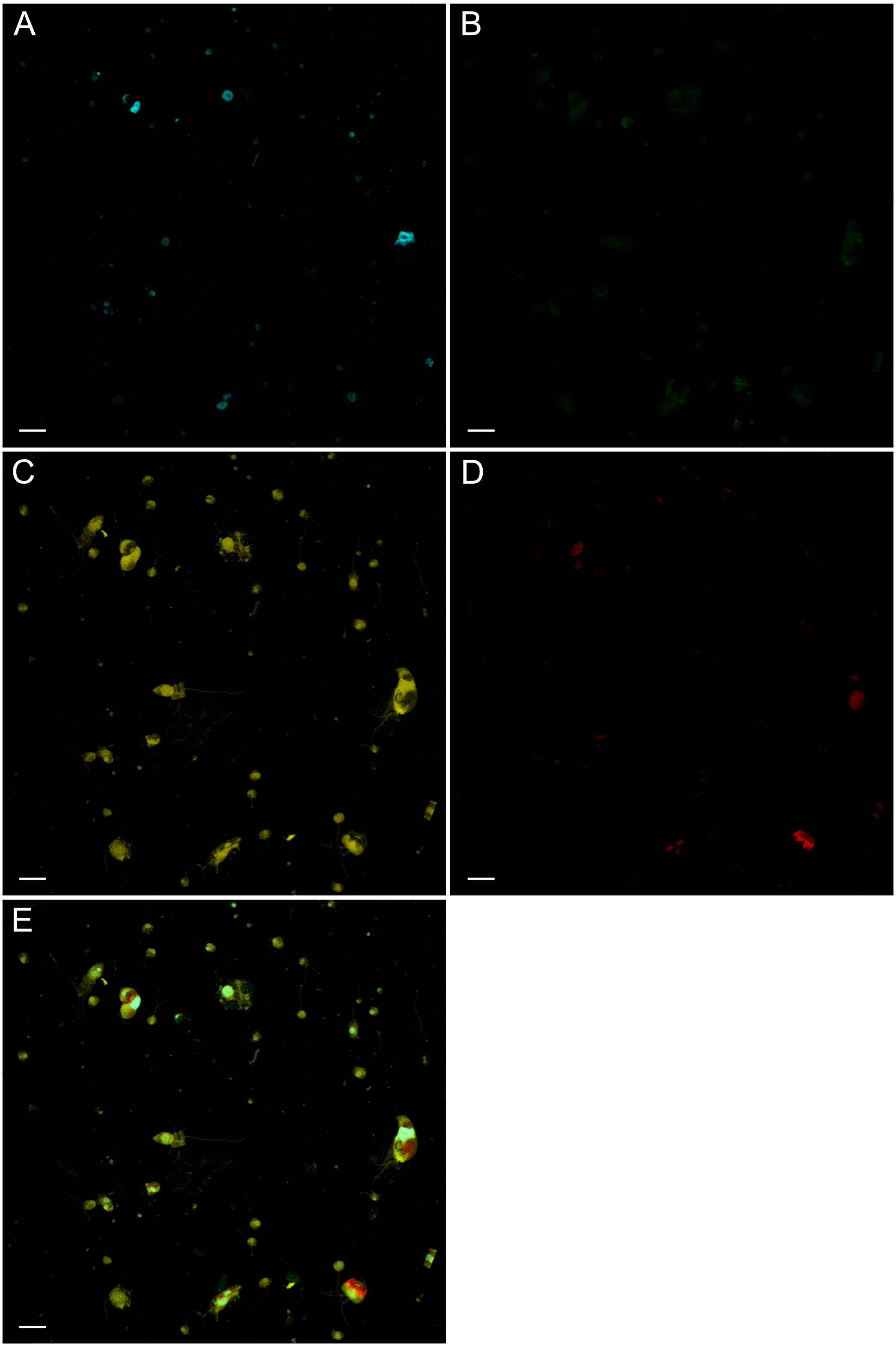
A
B
C
D
E

**Figure S13** Overview of filter from the Bellingshausen Strait sample serving as control for non-specific binding of the DEEPOL2 probe. (**A**) shows DAPI staining of nucleic acids, (**B**) shows no hybridization to non-target cells under the presence DEEPOL2-specific CARD-FISH probe, (**C**) NHS-ester protein staining using DY-560, (**D**) shows chlorophyll autofluorescence and (**E**) an overlay of all channels. Images shown represent maximum-intensity projections of Z-stacks. The scale bars correspond to 10 µm.

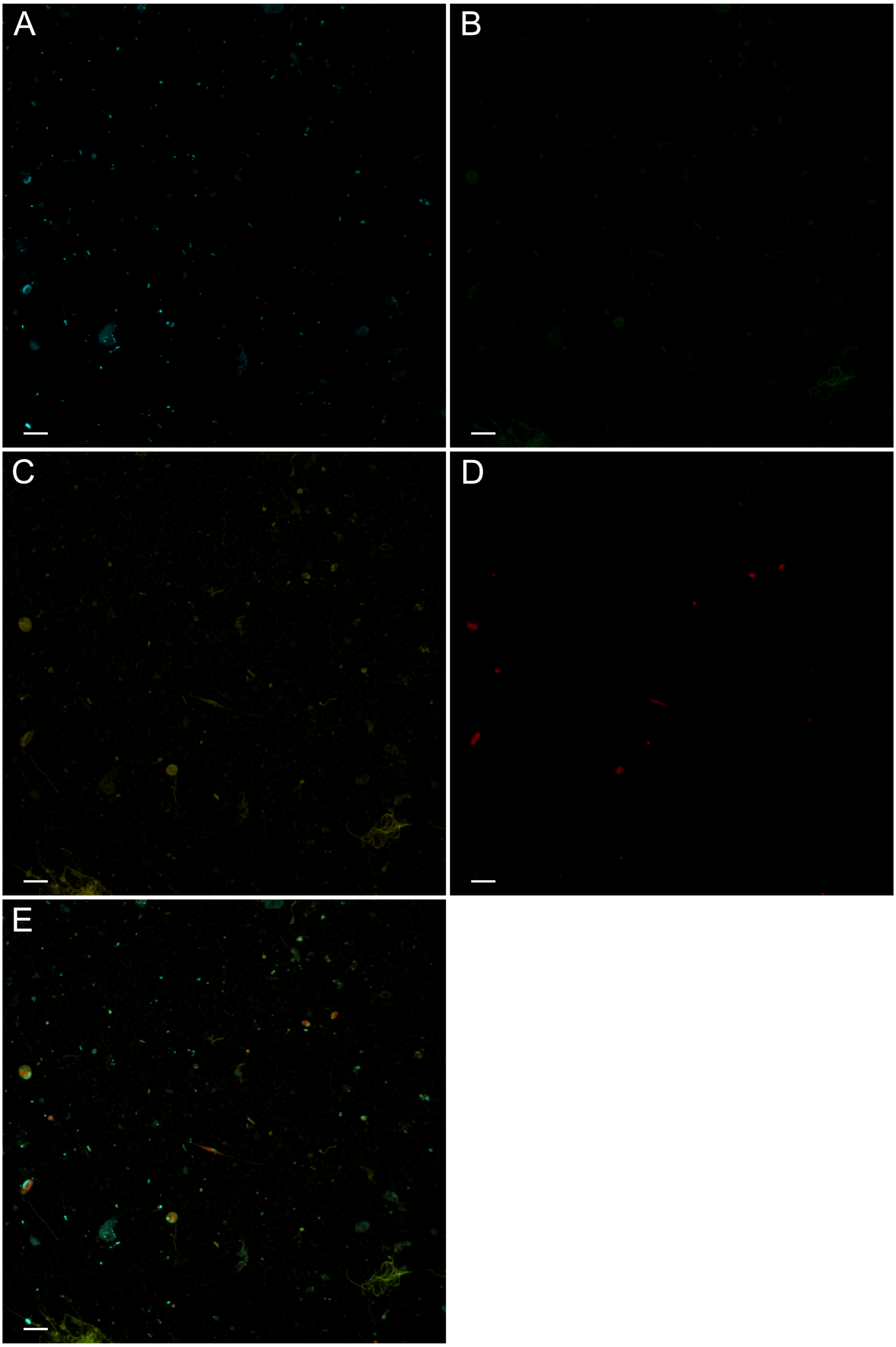
A
B
C
D
E

**Figure S14** Negative control applied to the sample KLM from the Skagerrak region. Panel (**A**) shows DAPI staining of nucleic acids, (**B**) shows no detected probe signal, (**C**) shows NHS-ester DY-560 protein staining, (**D**) shows chlorophyll autofluorescence and (**E**) an overlays of all channels. Images shown represent maximum-intensity projections of Z-stacks. The scale bars correspond to 10 µm.

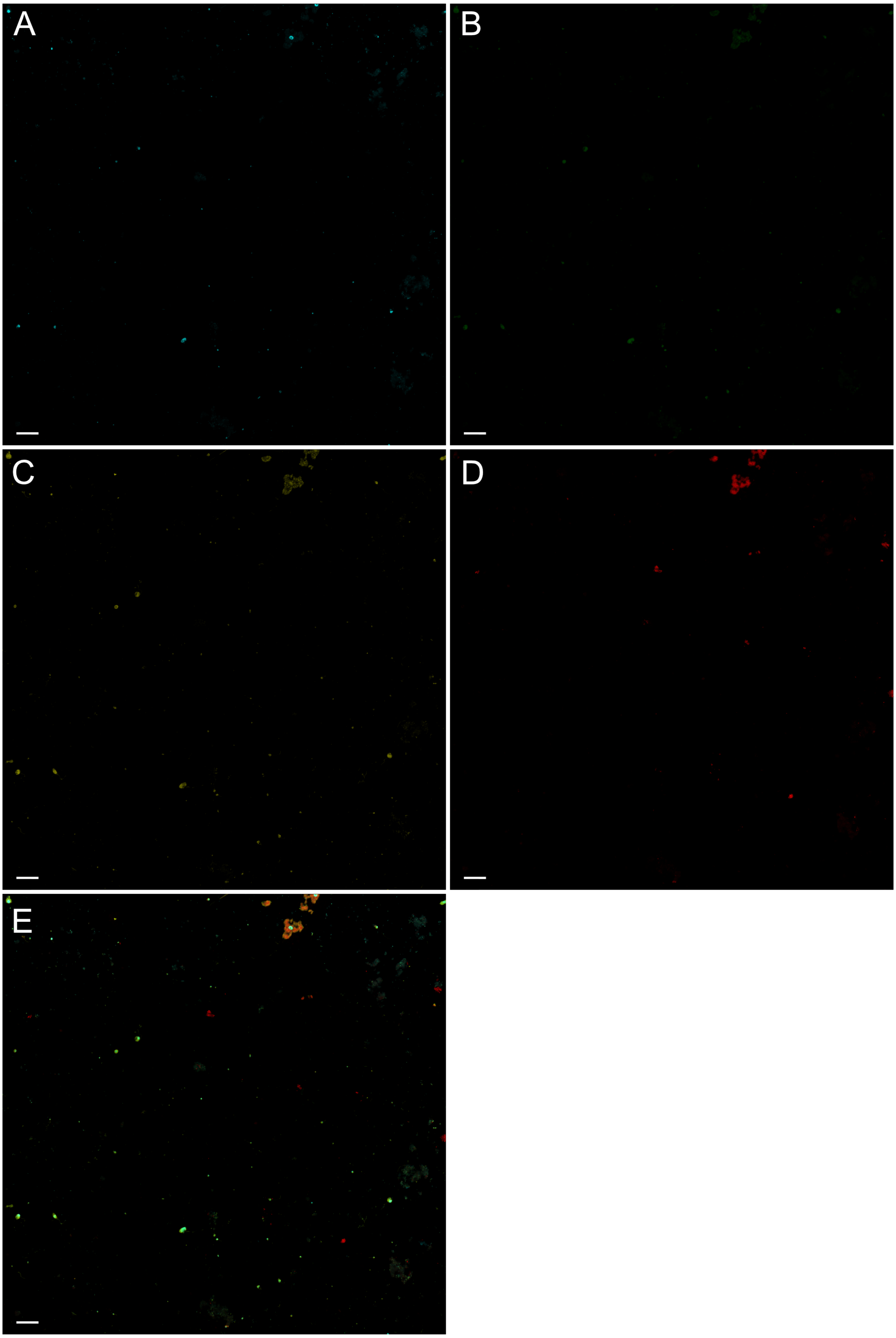
A
B
C
D
E

**Figure S15** Negative control applied to the sample KLD from the Skagerrak region. Panel (**A**) shows DAPI staining of nucleic acids, (**B**) shows no detected probe signal, (**C**) shows NHS-ester DY-560 protein staining, (**D**) shows chlorophyll autofluorescence and (**E**) an overlays of all channels. Images shown represent maximum-intensity projections of Z-stacks. The scale bars correspond to 25 µm.
